# Developmental regulation of intestinal *best4*+ cells

**DOI:** 10.64898/2025.12.17.694935

**Authors:** Abhinav Sur, Ella X. Segal, Michael P. Nunneley, Jason W. Sinclair, Morgan Kathleen Prochaska, Louis E. Dye, Yalan Wu, Liezhen Fu, Yun-Bo Shi, James Iben, Benjamin Feldman, Jeffrey A. Farrell

## Abstract

*best4*+/CFTR-high expressing cells are a recently described intestinal epithelial cell type potentially altered in inflammatory bowel disease and colorectal cancer. However, their developmental origin, developmental regulation, and functions remain undefined. This study identifies their conserved transcriptional program and uses zebrafish to dissect their developmental regulation *in vivo*. Lineage tracing identified that *best4*+ cells arise from *atoh1b*+ secretory progenitors. We identify that Notch signaling, mediated by *dll4,* specifies *best4*+ cells at the expense of enterochromaffin cells. Downstream of Notch, *meis1b* confers *best4*+ cell identity. *best4*+ cells then exhibit regionalized gene expression, regulated by *pbx3a*. Additionally, this study demonstrates a system where *best4*+ cells can be manipulated, observed, and removed in an organismal context. Live imaging and electron microscopy of *best4*+ cells identified dynamic cellular projections, suggesting a sensory or communicative function. Removal of *best4*+ cells *in vivo* eliminated previously proposed functions: they are not required to restore intestinal pH following acidic challenge and do not absorb nutrients. However, we identify region-specific intracellular pH differences that suggest potential functional heterogeneity. Altogether, this study presents a comprehensive description of *best4*+ cell development from birth to spatial regulation that will be instrumental to understand how *best4*+ cells change in disease or might be therapeutically manipulated and presents the tools to dissect their function *in vivo*.

## Introduction

To support its primary role of nutrient absorption, the gastrointestinal tract performs several additional critical functions to regulate metabolism, maintain fluid and electrolyte homeostasis, form a protective mucosal barrier, and respond to pathogenic threats. These functions are performed by distinct specialized intestinal epithelial cell types that are continuously replaced throughout an organism’s lifespan. While most intestinal epithelial cells (IECs) are well documented, we know relatively little about *best4+/*CFTR-high-expressing cells^1–5^. CFTR-high-expressing cells were first identified in rats via CFTR (Cystic Fibrosis Transmembrane conductance Regulator) immunohistochemistry in 1995^5^ and were first molecularly described as *best4*+ cells in the human colon in 2019 using single-cell RNA sequencing (scRNAseq)^2,3^. Additional scRNAseq studies have now identified *best4*+ cells in several species, including cows^6^, yaks^7^, pigs^8–11^, rabbits^12^, rats^5,13,14^, pythons^15^, *Xenopus*, zebrafish^16–18^, and sturgeon^19^, suggesting these cells are broadly conserved. *Drosophila*^20,21^ and mosquito^22^ scRNAseq did not identify *best4*+ cells, suggesting that they are a vertebrate-specific intestinal cell type. Interestingly, *best4*+ cells are absent from mice, likely due to evolutionary loss^23–25^. Though the functions of *best4*+ cells remain unknown, their gene expression profile has suggested potential involvement in fluid secretion, pH sensing or regulation, and immune response^16^. scRNAseq studies have also shown alteration in *best4*+ cells in gastrointestinal disorders like inflammatory bowel diseases^2,3^ and potentially colorectal cancer^26,27^, though whether there is any causative effect remains unknown. The broad evolutionary conservation of *best4*+ cells suggests they perform a crucial function and necessitates investigation into their contribution to intestinal homeostasis.

scRNAseq efforts from our lab and others have captured *best4*+ cells in zebrafish larval^16–18^ and adult stages^28^, and we previously demonstrated that zebrafish *best4*+ cells are transcriptionally similar to those in humans^16^. Because *best4*+ cells are absent from mice, one of the most popular experimental models, we propose that zebrafish provide an excellent opportunity to study these cells *in vivo,* in the context of surrounding signals and tissues in an organism with excellent genetic tractability, experimental accessibility, high fecundity, and optical transparency. Additionally, larval zebrafish development provides a unique opportunity to study the *de novo* formation of the intestine, when cell type specification is relatively synchronized. Zebrafish develop a functional intestine over 5 days, where Notch signaling first specifies intestinal progenitors into absorptive and secretory fates (30–70 hours) which then further differentiate into multiple subtypes of enterocytes, enteroendocrine cells (EECs), goblet cells, *best4*+ cells, and tuft-like cells that resemble those in mammals^16,17,29–34^ (Fig. 1A).

**Figure 1:**
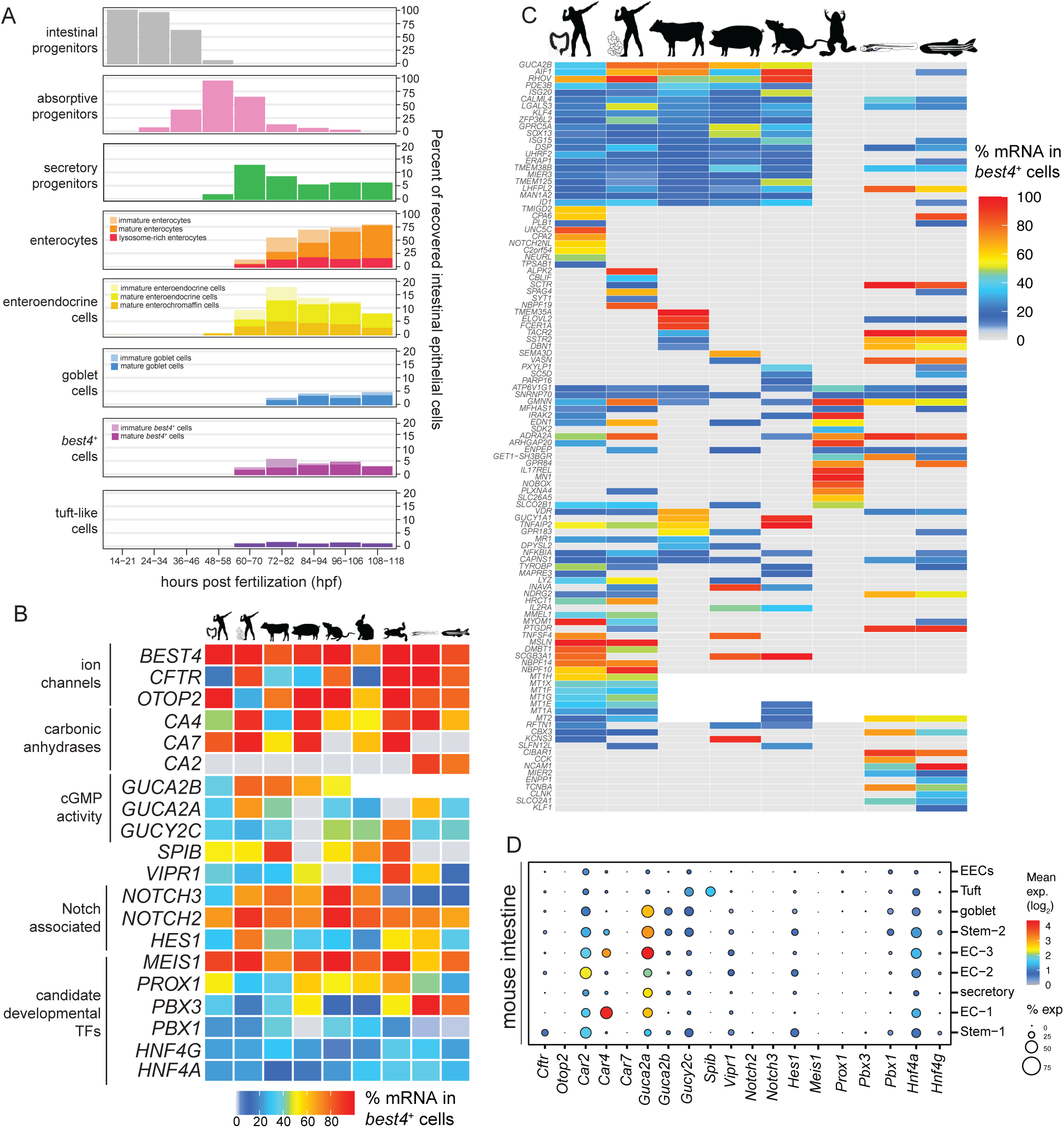
*best4*+ cells share a core conserved gene expression program across species. (**A**) Emergence of intestinal epithelial cell (IEC) types and subtypes during zebrafish larval development shown as percentage of IECs per stage in Daniocell (Sur et al. 2023). (**B–C**) Specificity of genes enriched in *best4*+ cells in (**B**) most profiled animals or (**C**) some profiled animals. Color represents percentage of mRNA within *best4*+ cells: average expression within *best4*+ cell cluster, divided by sum of average expression in all IEC clusters. Grey represents no expression, while white represents an unannotated gene. (**D**) Expression of conserved *best4*+ cell-specific genes in adult mouse colonic IECs (Davidi et al., 2025). See also Figure S1.

Given their recent molecular description, almost nothing is known about the origins, developmental regulation, or function of *best4*+ cells. Several studies have made computational predictions about the development of these cells using trajectory analysis^2–4,14,16^, but few of these predictions have been experimentally tested. Additionally, no *in vivo* system has been developed that allows observing *best4*+ cells in their native context or removing them to explore their function. Finally, almost no elements of their gene regulatory network have been identified thus far.

Using scRNAseq developmental trajectory analysis, we previously predicted that *best4*+ cells descend from secretory progenitors, are specified by Notch signaling, and that *dacha*, *meis1b*, and *pbx3a* may be key transcription factors (TFs) important for their specification^16^. In this study, we develop novel tools to study *best4*+ cells, including transgenic lines to perform live imaging and genetically remove *best4*+ cells *in vivo* for the first time. We test our previous developmental predictions by (1) lineage tracing *best4*+ cells, (2) testing the role of Notch signaling in their specification, and (3) combining new whole-gene deletion TF mutants with scRNAseq to determine the roles of *meis1b* and *pbx3a* in *best4*+ cell development. We additionally explore the *in vivo* effects of loss of *best4*+ cells for the first time.

## Results

### Cross-species analysis of *best4*+ cell gene expression similarity

Since recent intestinal scRNAseq datasets have identified *best4*+ cells in diverse species, including mammals (human, cow, pigs, rats, rabbits) and aquatic vertebrates (*Xenopus*, zebrafish), we sought to identify the core gene expression program shared across *best4*+ cells in all animals. Thus, we identified genes expressed specifically within *best4*+ cells among IECs across several published scRNAseq datasets (Fig. 1B–C, Fig. S1A–B). This revealed a core set of functional genes that were universally shared, including *best4*, *cftr*, *otop2*, *carbonic anhydrases (ca2, ca4, ca7)*, guanylin (*guca2a*) and uroguanylin (*guca2b*) (Fig. 1B, Fig. S1A). As others have proposed^1–3,12,35,36^, the presence of these genes indicate that key conserved functions of *best4*+ cells may include bicarbonate ion conversion and transport (*best4, cftr, ca2/4/7*), fluid/electrolyte secretion (*cftr, guca2a/2b, gucy2c*), proton influx or pH homeostasis (*otop2, ca2/4/7*), cGMP activity (*guca2a/2b*, *gucy2c*), and response to vasoactive intestinal peptide (*vipr1*) (Fig. 1B, Table 1). In some cases, functions appear to be conserved, despite different genes encoding those functions. For example, among carbonic anhydrases, zebrafish express *ca2*, rather than *ca7*, but these genes seem to have similar functions and localization^37^ (Fig. 1B). Additionally, while hormone annotations are potentially incomplete in zebrafish, we only detect *guca2a* (guanylin) expression, while mammalian *best4*+ cells express both *guca2a* (guanylin) and *guca2b* (uroguanylin), though these hormones target the same guanylate cyclase (*gucy2c*)^38–40^.

**Table 1:**
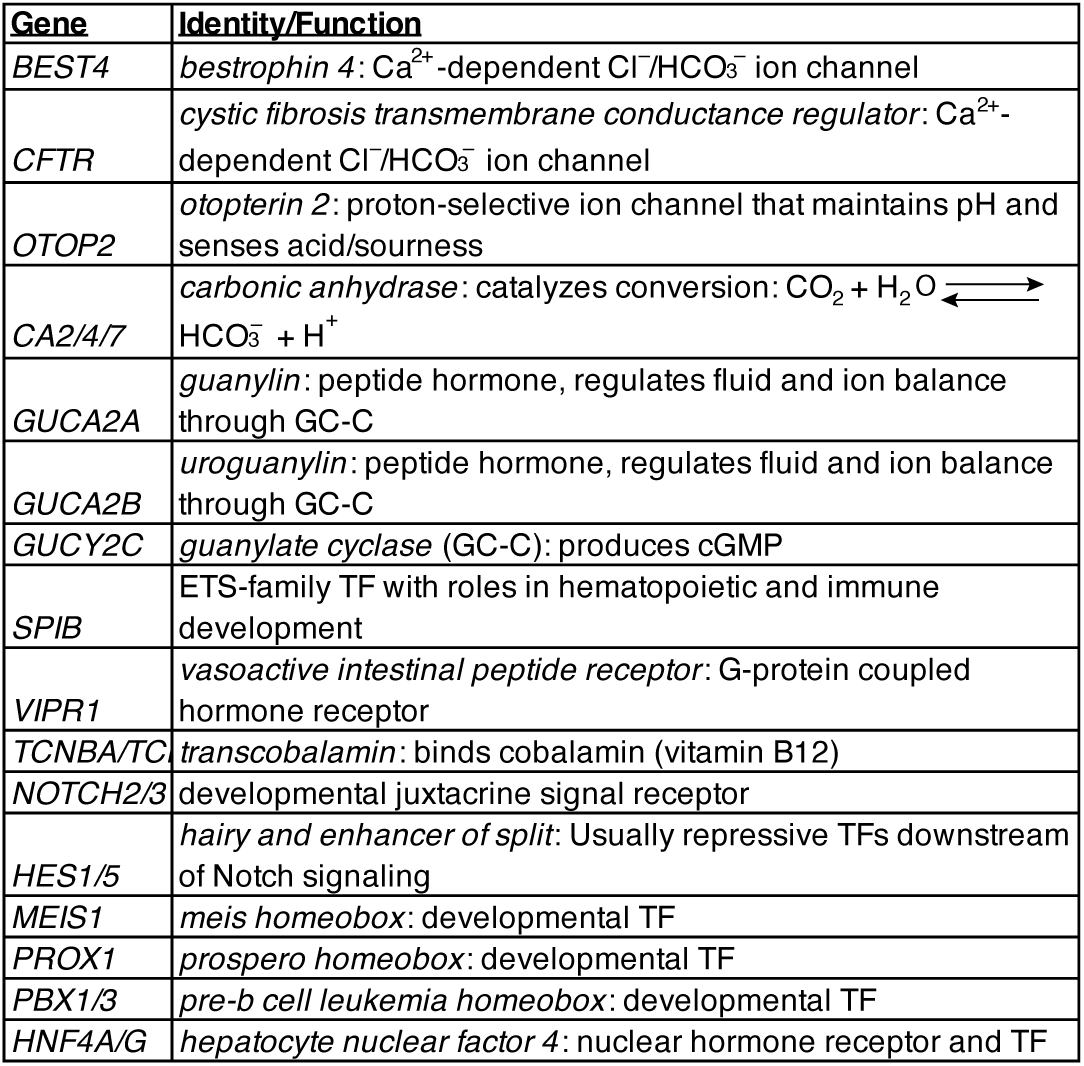
Known details about conserved *best4+*-cell specific marker genes.

In addition to core, shared functional genes, we observed several developmental regulators that are expressed in *best4*+ cells across all species, suggesting that the developmental program that specifies these cells is also conserved. First, in terms of developmental signals, all *best4*+ cells express Notch2 and Hes1 orthologs, which are typically Notch targets (Fig. 1B). Additionally, all *best4*+ cells express a group of cell type-specific transcription factors, including *MEIS1*, *PROX1*, and *PBX1/3*. Interestingly *SPIB*, which was recently proposed as an important regulator of *best4*+ development^36,41^, was expressed in *best4*+ cells in many animals, but not observed in pigs or zebrafish (Fig. 1B). Altogether, this analysis highlights the developmental and functional programs that are shared in *best4*+ cells across evolution and proposes key targets to test in animal models to better inform our understanding of *best4*+ cells across species.

We also observed groups of genes or putative functions (Fig. 1C; Fig. S1B) that were specific to *best4*+ cells in several species, but less completely conserved, indicating either evolutionary variation, potential convergent evolution, or possibly limitations of this analysis (e.g. ortholog annotation and mapping, differences in developmental stage, or particular scRNAseq datasets) (Fig. 1C). For instance, some developmental signals were expanded: mammalian *best4*+ cells also express *NOTCH3* (Fig. 1B), and human colonic *best4*+ cells additionally express *Notch2 N-terminal-like* (*NOTCH2NL*) (Fig. 1C). Also, some less broadly shared TFs were observed, including the previously mentioned *SPIB*, BMP-responsive *ID1*, and *CBX3* (Fig. 1C). Several neuroactive G-protein coupled hormone receptors were conserved between zebrafish and some, but not all, other species (Fig. 1C). For instance, the alpha-adrenergic receptor (*adra2a*) is expressed in human, rat, *Xenopus*, and zebrafish *best4*+ cells; secretin receptor (*sctr)* and prostaglandin receptor (*ptger4c)* are shared between human small intestinal and zebrafish *best4*+ cells; and tachykinin receptor (*tacr2*) and somatostatin receptor (*sstr2*) are shared between cow and zebrafish *best4*+ cells (Fig. 1C; Fig. S1B). This suggests that *best4*+ cells are highly communicative with signaling partners either within or outside the intestine, potentially responding to adrenergic signaling (*adra2a*), luminal nutrients (*sctr*), pH (*sctr*), intestinal inflammation (*ptger4c*), or other aspects of the intestinal environment. *best4*+ cells in multiple species also specifically express machinery involved in vitamin B12 absorption^42,43^ — *best4*+ cells in human small intestines express the *cobalamin binding intrinsic factor* (*CBLIF*) and zebrafish *best4*+ cells express *transcobalamin* (*tcnba*) (Fig. 1C, Fig. S1B–C). Finally, *best4*+ cells specifically express metallothioneins (MT) in zebrafish (*mt2*), rats (*mt2, mt1a, and mt1e*), and humans (*mt1e*, *mt1f, mt1g*, *mt1h*, *mt1a*, *mt1x*, and *mt2*) (Fig. 1C). Metallothionein expression may enable *best4*+ cells to protect against toxic metal consumption or to uptake zinc ions, a crucial co-factor of carbonic anhydrases, which are enriched in *best4*+ cells^44–48^. Altogether, these examples suggest that there may be more functional similarity between *best4*+ cells in different animals than suggested by universally expressed genes, including detoxification, communication with other cells in the digestive system, inflammatory response, and vitamin B12 transport.

Since mice have lost *best4*+ cells, it raises the question of whether *best4*+ cell marker genes are expressed elsewhere in mouse intestines. Thus, we identified which intestinal cell types in C57BL/ 6J post-weaning mice express conserved *best4*+ cell-specific genes using previously published scRNAseq data^49^. *best4*+ cell-specific developmental regulators (*Notch2*, *Notch3*, *Prox1*, *Meis1*, *Pbx3*) were not expressed in any mouse IECs (Fig. 1D). Also, a few conserved *best4*+ cell functional genes were absent from the mouse intestine. For example, *Best4* is a pseudogene and *Otop2* is not expressed, indicating that the mouse intestine does not perform some *best4*+-cell functions. However, most *best4*+ cell functional genes are expressed across all mouse IECs (e.g., *Car2*, *Guca2a*, *Guca2b*, *Cftr, Vipr1*), suggesting that most of their functions persist, but the regulation of those functions may be different in mice. Furthermore, the absence of the conserved developmental program in an animal where *best4*+ cells are absent further suggests these factors will be instrumental in *best4*+ cell specification.

### *best4*+ cells exhibit dynamic morphologies with vesicle-laden protrusions

Given the number of cell-surface ion channels and receptors specific to *best4*+ cells across animals, we next investigated whether *best4*+ cells exhibited unique morphological features that might suggest interaction with their environment or neighbors. The optical transparency of zebrafish larvae provides a unique opportunity to study and image *best4*+ cells live in their full tissue context (Fig. 2A). To do so, we generated Tg(*best4:eGFP, cryaa:mCherry*)^y7^^27^ and Tg(*best4:mApple, cryaa:eGFP*)^y726^ transgenic fish using a 2.3 kb region ∼4.1 kb upstream of *best4* driving expression of GFP or mApple, randomly inserted in the genome using Tol2 transposase (Fig. 2B, S2A), which accurately labeled *best4*+ cells in the intestine (Fig. S2B–B’). Timelapse imaging of Tg(*best4*:*mApple*)^y726^ intestines revealed a diversity of *best4*+ cell shapes and that *best4*+ cells dynamically extend and retract cellular protrusions over a timespan of multiple hours (Fig. 2B–D, yellow and cyan arrowheads, Video S1). Some *best4*+ cells mirrored classical EEC cell shape^50,51^, with the cell body located away from the lumen (labeled with dextran) and a thin apical projection extending between neighboring epithelial cells toward the lumen (Fig. 2D, yellow arrowheads); other cells were columnar and touched the lumen (Fig. 2D, orange arrowheads), and sometimes extended basolateral projections, potentially toward adjacent epithelial cells, the intestinal stroma, or the enteric nervous system (Fig. 2D, cyan arrowheads). Projections were observed in *best4*+ cells throughout the length of the gut, suggesting that these are not region-specific cellular features (Fig. S2C).

**Figure 2:**
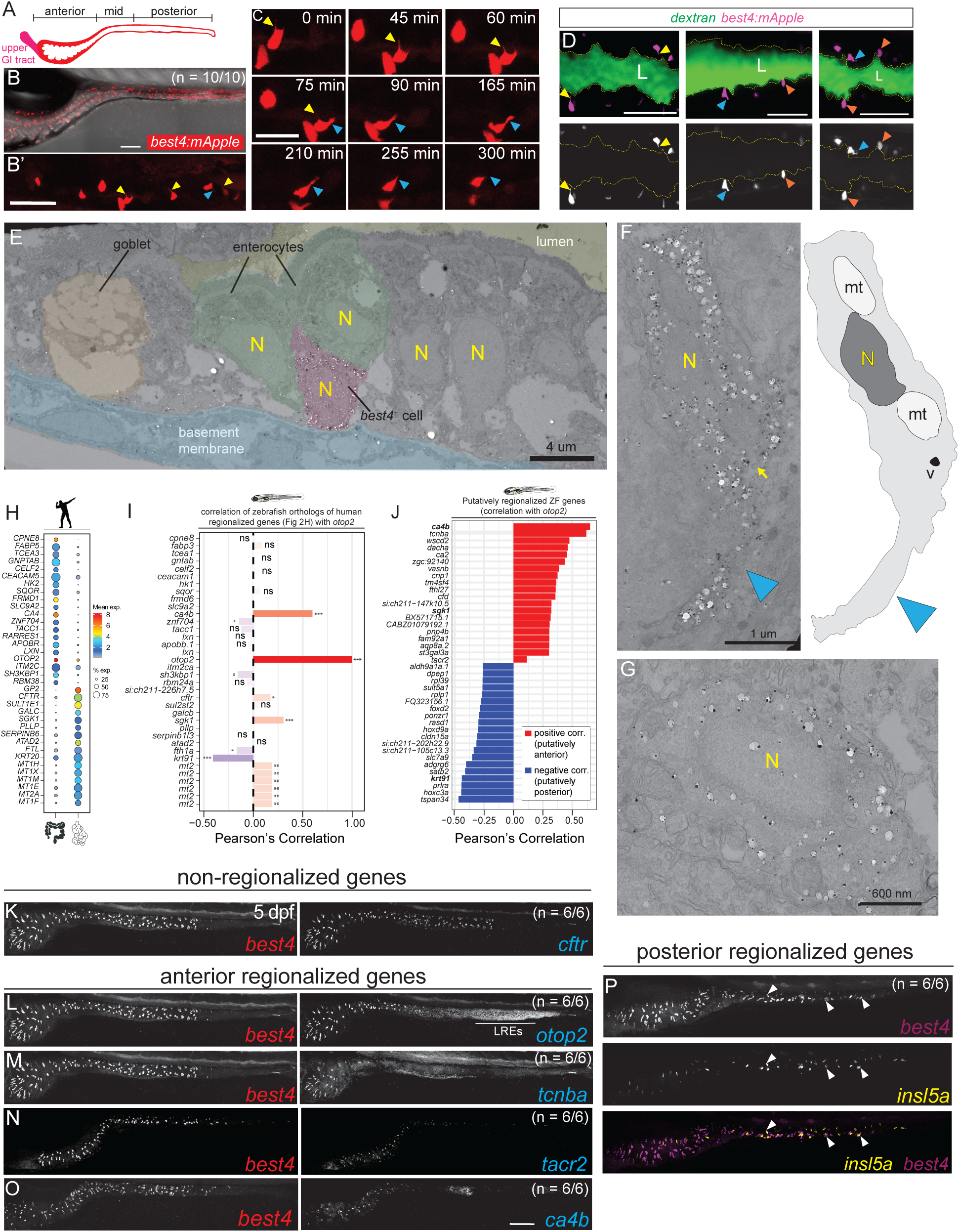
*best4*+ cells extend dynamic protrusions and are spatially heterogeneous. (**A**) Cartoon showing anterior, mid, and posterior intestinal regions. (**B–D**) Live imaging of *best4*+ cells in *Tg(best4:mApple)* 5 dpf zebrafish larvae in (**B**) the entire intestine, (**B’**) the intestinal bulb, (**C**) over time, and (**D**) in context of the intestinal lumen labeled with Alexa647-Dextran (“L”). Arrowheads indicate *best4*+ cells with apical projections (yellow), basolateral projections (cyan), or no projections (orange). Scale bars: 100 um (**B**), 50 um (**B’–D**). (**Legend continues on next page.**) (**E–G**) Immuno-electron microscopy in *Tg(best4:eGFP)* 7 dpf larvae. Black dots with white halos are silver-enhanced anti-GFP immuno-gold. (**E**) *best4*+ cells in the context of the intestinal epithelium and (**F**) higher magnification view of individual *best4*+ cells with a cytoplasmic protrusion (cyan arrowhead) or (**G**) without a protrusion. Yellow arrow denotes a vesicle. “N”: nucleus, mt: mitochondria, v: vesicle. (**H–I**) (**H**) Differentially expressed genes between human colonic and small intestinal *best4*^+^ cells and (**I**) the correlation of their zebrafish orthologs with *otop2,* an anteriorly regionalized *best4*+ cell marker. *p* < 0.05 (*), 0.01 (**) or 0.01 (***); ns: not significant. (**J**) Zebrafish *best4*+ cell-enriched genes correlated and anti-correlated with *otop2*. (**K–P**) HCR of *best4*+ cell markers that are (**K**) not regionalized, (**L–O**) anteriorly regionalized, or (**P**) mid/ posterior regionalized. *otop2* is also expressed in lysosome-rich enterocytes (LREs). Scale bar: 100 um. See also Figures S2 and S3.

We also used *Tg(best4:eGFP)* ^y727^ to investigate *best4*+ cell subcellular architecture using immuno-electron microscopy (EM). A gold-tagged anti-GFP antibody was used to identify cytoplasmic GFP within *best4*+ cells in serially sectioned 7 dpf *Tg(best4:eGFP)* ^y727^ zebrafish larvae (Fig. 2E–G, Fig. S2D–F). As observed in live imaging, some *best4*+ cells had a classical EEC shape^51^, tucked away from the lumen between enterocytes (Fig. 2E). However, the most distinctive *best4*+ cells were those with basolateral cytoplasmic projections (Fig. 2F, Fig. S2D–F, cyan arrowheads). In EM, it was apparent that vesicles (identified as membrane-bound structures that were heterogenous in size with empty lumens) were sometimes present at the base of those projections (Fig. 2F, Fig. S2E, F, yellow arrow) and occasionally within the projections themselves (Fig. S2D, arrow). *best4*+ cells were also observed without cytoplasmic projections (Fig. 2G), consistent with their dynamic nature, as observed during live imaging (Fig. 2B–C). Beyond these cytoplasmic projections, no additional remarkable subcellular features were observed in *best4*+ cells, consistent with the fact that they were not initially morphologically identified in historical EM studies without labeling^52–56^. These projections may be a conserved feature, since immunohistochemistry against BEST4 shows basolateral processes in rats^13^ and potentially human organoids^36^. This study identifies a diversity and dynamic nature of *best4*+ cell shapes, multiple orientations of protrusions, and vesicles within some protrusions, which suggest that *best4*+ cells likely act to integrate sensory information or signal to other cells, and potentially have multiple signaling partners over time.

### *best4*+ cells exhibit spatial transcriptional heterogeneity

In humans, *best4*+ cell gene expression varies in different regions of the gut: when comparing the small intestine and colon, CFTR is expressed only in small intestinal *best4*+ cells, while the proton channel OTOP2 is expressed more strongly in colonic *best4*+ cells than small intestinal *best4*+ cells, where its expression is ileal-specific (Fig. S1A)^1–4^. We previously demonstrated that zebrafish *best4*+ cells also exhibit regionalized gene expression^16^, with *otop2* highly regionalized, but to a potentially different region: anterior *best4*+ cells. We searched published human scRNAseq data to more broadly identify *best4*+ cell markers that were differentially expressed between the colon and small intestine (Fig. 2H) and calculated their correlation to *otop2* as a measure of spatial regulation in zebrafish (Fig. 2I). Only a few genes were spatially regulated in both organisms (*ca4*, *otop2*, *sgk1*, and *krt91/KRT20*), suggesting that spatial regulation may be less conserved between species. Unlike in humans, *cftr* is not spatially regulated in zebrafish larvae (Fig. 2I). However, to identify other spatially heterogenous genes within zebrafish *best4*+ cells, we identified genes in our scRNAseq data whose expression was strongly correlated or anti-correlated within *best4*+ cells to the anterior marker *otop2* (Fig. 2J). This identified several genes that are also restricted to anterior *best4*+ cells, including *ca4b (*carbonic anhydrase), *tcnba* (a vitamin B12 transporter), *wscd2* (an O-sulfotransferase), *dacha* (a transcription factor), *ca2*, *vasnb*, *cfd,* and *tacr2* (tachykinin receptor 2), among others. Additionally, several genes had anti-correlated expression, suggesting they are restricted to posterior *best4*+ cells (Fig. 2J). Some regionalized genes were very specific to *best4*+ cells within the gut, including *adgrg6* (a steroid hormone receptor) and *sult5a1* (a sulfotransferase), while others were expressed in several posterior intestinal cell types, including *tspan34*, *prlra*, *slc7a9*, *foxd2*, and *dpep1*.

We then validated the spatial expression patterns of putatively spatially regulated *best4*+ cell markers (Fig. 2K–P, Fig. S3A–L) using HCR (Hybridization Chain Reaction RNA *in situ* hybridization). These imaging data confirmed that *cftr* is expressed in all zebrafish *best4*+ cells (Fig. 2K, Fig. S3B, E, H, K) and that *otop2, tacr2*, *tcnba*, and *ca4b* are restricted to anterior *best4*+ cells (Fig. 2L–O, Fig. S3D, G, J, F, I, L). Interestingly, the anterior-posterior expression domain varied for each gene, suggesting that each gene’s regulatory elements are independently interpreting spatial context. Since gut development proceeds from anterior to posterior, we tested whether this spatial heterogeneity might simply reflect developmental asynchrony by assessing regionalized expression across several developmental stages (3–7 dpf) (Fig. 2K–O; Fig. S3A–L). The expression domains of *cftr*, *otop2*, and *tcnba* remained constant across stages indicating an intrinsic, temporally consistent transcriptional heterogeneity. Additionally, via staining, we unintentionally identified that *insl5a* is expressed only within mid-posterior *best4*+ cells, in addition to EECs (Fig. 2P), though we did not identify similarly expressed genes. Altogether, these results suggest that *best4*+ cells have stable spatial heterogeneity, and their spatially restricted markers may indicate different functions for these cells in different regions of the gut, but this regionalization may differ between species.

### *best4*+ cells originate from *atoh1b*+ secretory progenitors

Since many developmental factors in *best4*+ cells are shared across most animals (Fig. 1B), zebrafish provide an excellent model to investigate their development *in vivo,* without the need for exogenous application of developmental factors. One key topic of debate is whether *best4*+ cells arise from absorptive or secretory progenitors, since they express genes characteristic of both absorptive and secretory cells^3,4,12,14,16^, and computational analyses have linked them to both progenitor types across different systems^2–4,14,16^. Based on our previous scRNAseq transcriptional trajectory (Fig. 3A), we suggested that *best4*+ cells arise from secretory progenitors, and we aimed to experimentally validate this prediction. Secretory progenitors are characterized by expression of *ascl1a*, *atoh1b*, *tnfrsf11a*, *sox4a*, and *sox4b*^16^ (Fig. S4A), but these progenitor transcripts were not observed in *best4*+ cells in our scRNAseq data. We first used HCR as a more sensitive technique to search for co-expression that might tie *best4*+ cells to secretory progenitors. However, co-expression of *best4* mRNA was not observed with *atoh1b* (Fig. S4B), *ascl1a* (Fig. S4C), *tnfrsf11a* (Fig. S4D), *sox4a* (Fig. S4E), or *sox4b* (Fig. S4F) in the zebrafish gut, suggesting these transcripts are effectively eliminated by the time *best4*+ cells differentiate.

**Figure 3:**
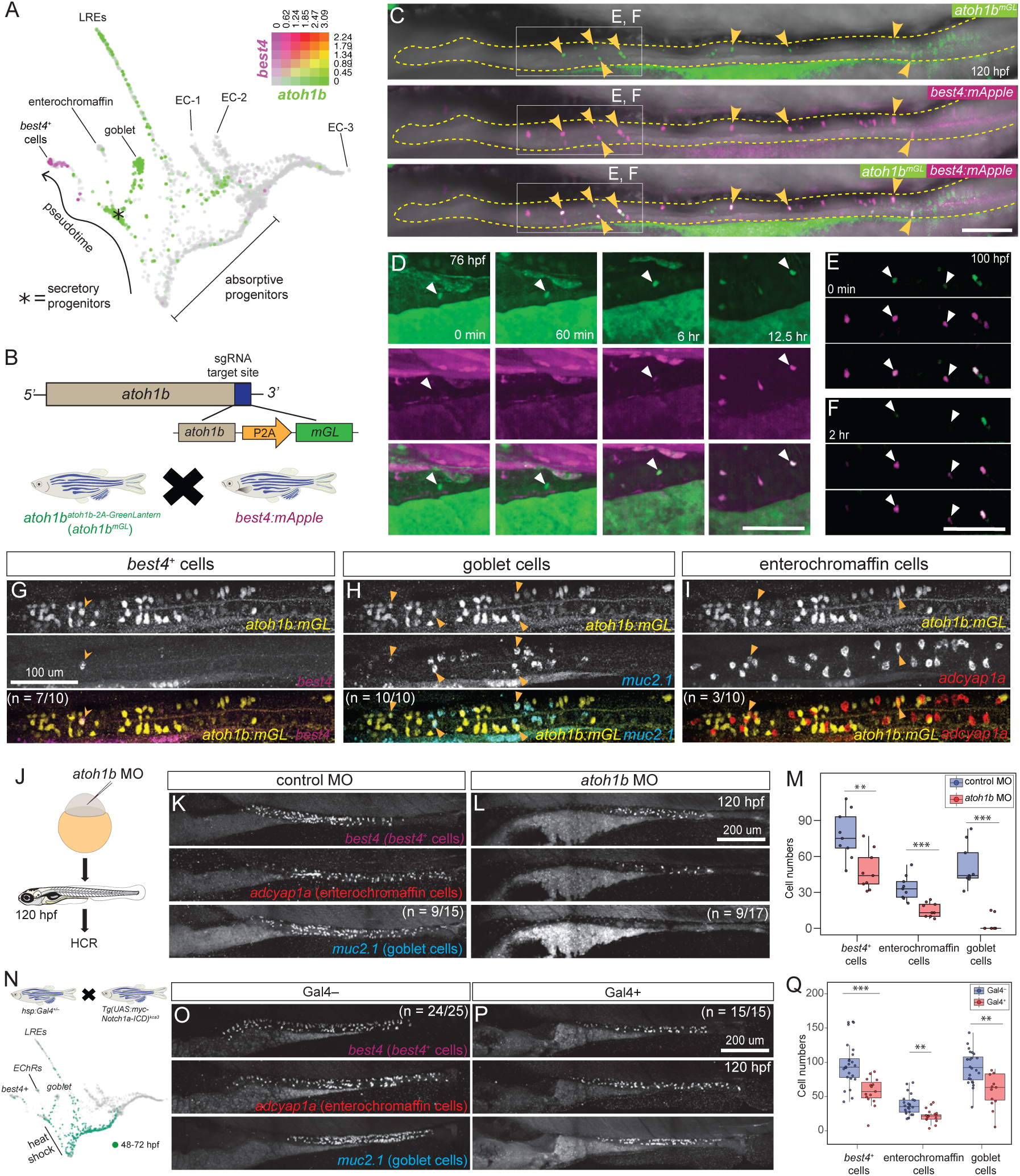
*best4*+ cells have a secretory origin. (**A**) Co-expression of *atoh1b* (green) and *best4* (magenta) on URD-inferred transcriptional trajectory of intestinal epithelial cells (Sur et al 2023). Co-expression matrix shows expression levels (log2) for each gene and resulting cell color in plot. (**B**) Schematic of *atoh1b^atoh1b:2A-mGreenLantern^* endogenous knock-in line. (**C–F**) (**C**) Live confocal micrographs of *atoh1b^mGL^*; Tg*(best4:mApple)*. Yellow arrowheads: mGL+/mApple+ cells. (**D**) Onset of mApple fluorescence in mGL+ secretory progenitor and (**E–F**) decline of mGL fluorescence in an mApple+ cell. Scale Bars: 100 um. **(G–I)** HCR and anti-GFP immunofluorescence in *atoh1b^mGL^* 4 dpf larvae marks mGL expression (arrowheads) in (**J**) *best4*+ cells, (**K**) enterochromaffin cells, and (**L**) goblet cells. (**J–M**) Reduction of secretory progenitors: HCR for *best4*+ cells (*best4*), enterochromaffin cells (*adcyap1a*), and goblet cells (*muc2.1*) after control MO or a*toh1b* MO injection. *p* < 0.01 (**) or 0.001 (***). (**N–Q**) Reduction of secretory progenitors: early Notch overexpression using Tg(*hsp:Gal4*); Tg(UAS:myc-Notch1a-ICD) fish, similar to J–M. See also Figures S4 and S5.

Thus, to follow secretory progenitors over time, we created an *atoh1b^atoh1b-2A-mGreenLantern^* endogenous knock-in line (*atoh1b*^y732^, hereafter *atoh1b^mGL^*) that labels secretory progenitors with the long-lived fluorescent protein mGreenLantern (mGL), since Atoh1 is often regarded as the master regulator of secretory fate^57–60^ (Fig. 3A–B). *atoh1b* mRNA is expressed in secretory progenitors and goblet cells, which also express *muc2.1* (Fig. 3A, Fig. S5A), and lysosome-rich enterocytes (LREs), which are specifically located in the posterior of the intestine. We validated the *atoh1b^mGL^* line using HCR and found that while goblet cells (*atoh1b^+^/muc2.1^+^*) were only sometimes labeled with mGL, secretory progenitors (*atoh1b^+^/muc2.1^−^*) were strongly labeled with mGL, and all mGL^+^ cells indeed expressed *atoh1b* mRNA (Fig. S5A), indicating that this line reliably marks secretory progenitors.

To determine whether *best4*+ cells arise from *atoh1b*^+^ secretory progenitors, we crossed Tg(*best4:mApple, cryaa:eGFP*)^y726^ with *atoh1b^mGL^* and performed live imaging (Fig. 3B). Cells marked by both *atoh1b^mGL^* and *Tg(best4:mApple)*^y726^ were observed along the entire anterior-posterior axis of the gut (Fig. 3C, Video S2). In all gut segments, some mGL+ cells progressively became mApple+ (Fig. 3D, white arrowhead, Video S3) between 72–88 hpf, shortly after they are first identifiable in scRNAseq (Fig. 1A). Additionally, mGL signal decayed from mGL+/mApple+ cells as they differentiated (Fig. 3E–F, white arrowhead, Video S2). We also observed several mGL+ progenitors that remained mApple–, suggesting they were persistently uncommitted secretory progenitors or adopted a different secretory fate (Video S2). Overall, these observations identified *best4*+ cells arising from *atoh1b*+ secretory progenitors in all regions of the zebrafish gut.

In mammals, recent studies have begun to uncover heterogeneity within secretory progenitors that may influence their downstream cell fates^36^. Thus, to determine which other secretory derivatives arise from *atoh1b*^+^ progenitors, we stained *atoh1b^mGL^* larvae for enterochromaffin cells (*adcyap1a^+^*), *best4*+ cells, goblet cells (*muc2.1^+^*), and tuft-like cells (*pou2f3*^+^) (Fig. 3G–I, Fig. S5H). The gut develops from the anterior to posterior, and mGL expression concordantly transitioned from the anterior to posterior between 4 and 5 dpf (Fig. S5B–G). Over these timepoints, we observed overlap of mGL^+^ cells frequently with *best4*+ cells and goblet cells (Fig. 3G–H, Fig. S5B–G), occasionally with enterochromaffin cells (at lower *atoh1b^mGL^* expression levels) (Fig. 3I), and never with tuft-like cells (Fig. S5H). An important caveat is that tuft-like cells are very rare at these stages, so their co-expression could have been missed. However, our results suggest that *best4*+ cells, goblet cells, and enterochromaffin cells share a common progenitor in zebrafish.

To further confirm whether *best4*+ cells arise from secretory progenitors, we performed multiple manipulations to alter secretory progenitor numbers and see whether these manipulations affected *best4*+ cell number. First, we knocked down *atoh1b* using a translation blocking morpholino (MO), which reduced enterochromaffin cells and *best4*+ cells and eliminated goblet cells (Fig. 3J–M). One potential reason for a stronger effect in goblet cells may be that *atoh1b* expression persists in them as they differentiate and may play a dual role in specification and differentiation (Fig. 3A). Second, since Notch signaling suppresses secretory fate during the absorptive versus secretory fate decision^30,57,61,62^, we reduced secretory progenitors by increasing Notch signaling (Fig. 3N–Q). Tg(*hsp:Gal4*); Tg(*UAS:myc-Notch1a-ICD*) were heat shocked from 48–72 hpf while secretory progenitor specification is occurring (Fig. 3N) to overexpress constitutively-active Notch, which significantly decreased *best4*+ cells, enterochromaffin cells (*adcyap1a*^+^), and goblet cells (*muc2.1*^+^) (Fig. 3O–Q). These results show that reduced *atoh1b* expression and leads to an overall reduction of secretory fate, and that reduction of secretory fates via multiple approaches reduces *best4*+ cell number. Altogether, we provide several lines of experimental evidence demonstrating that *best4*+ cells in multiple gut regions originate from secretory progenitors, particularly ones that express *atoh1b*.

### Notch signaling specifies *best4*+ cells among secretory progenitors

Since secretory progenitors give rise to several cell types, a key question is to understand which developmental signals specify secretory progenitors to become *best4*+ cells. Based on our previous developmental trajectory analysis (Fig. 4A, Fig. S6A), as well as conservation of *notch2* as a cell-type-specific marker of *best4*+ cells across evolution (Fig. 1B), we hypothesized that a second wave of Notch signaling within defined secretory progenitors is required for *best4*+ cell specification in zebrafish, potentially mediated by Notch2. We first tested which secretory derivatives had evidence of active Notch signaling at the time *best4*+ cells are specified using the Tg(*Tp1-bglob:eGFP*)^um14^ Notch reporter line^63^. At 120 hpf, reporter signal was exclusively observed in *best4*+ cells (Fig. 4B, Fig. S6C), suggesting that *best4*+ cells are the only Notch-responsive secretory cells. Indeed, scRNAseq data indicated that *best4*+ cells also specifically expressed *notch2* and *her15.1*/*her2* (Hes5) Notch signaling effectors (Fig. S6A–C). *best4*+/GFP^−^ cells were also observed (Fig. 4B, dotted circles), suggesting that *Tp1:GFP* expression (and thus Notch signaling) is transient in *best4*+ cells and likely not required for maintenance of their identity.

**Figure 4:**
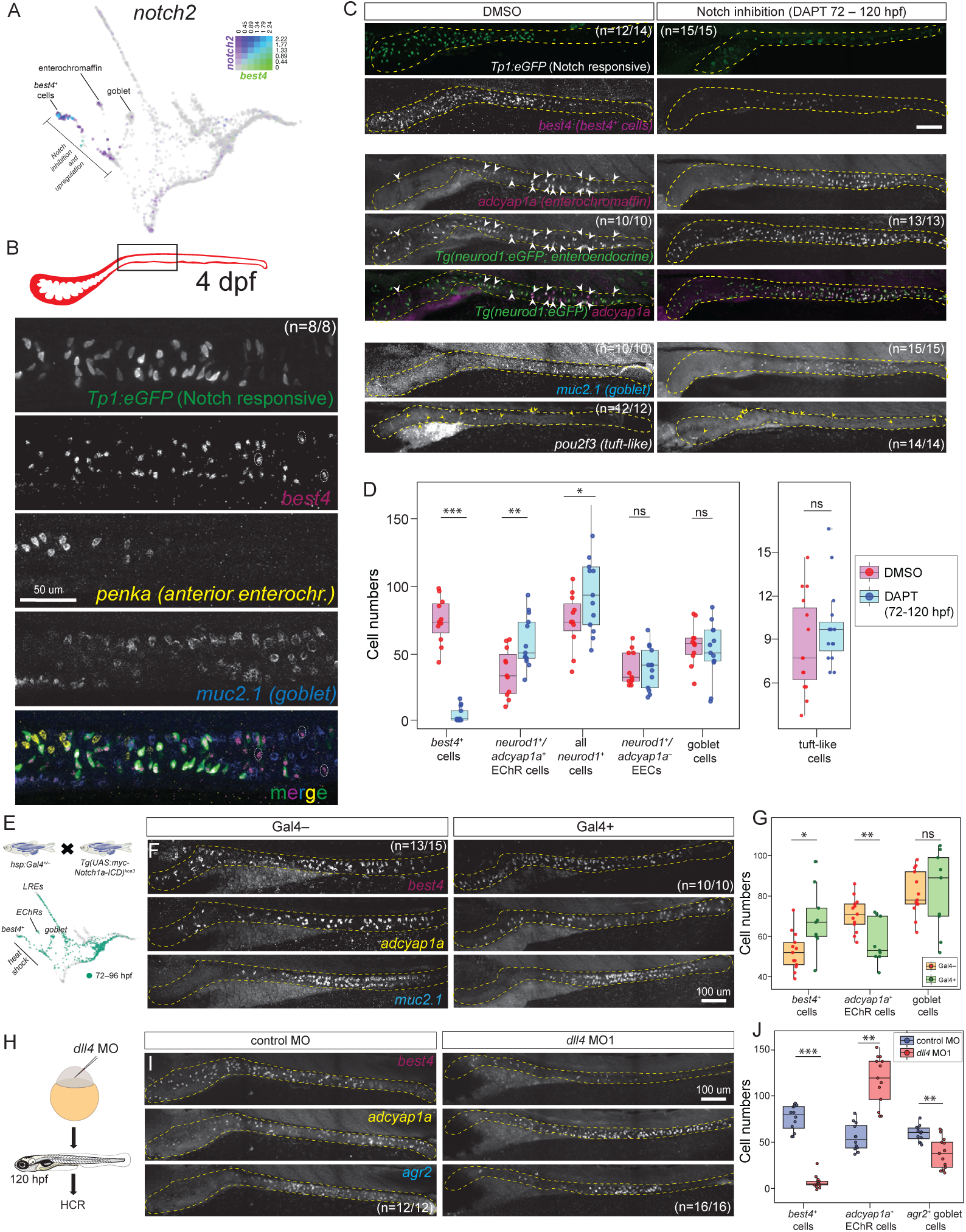
Notch signaling through *dll4* triggers *best4+* cell specification. (**A**) Co-expression of *notch2* (purple) and *best4* (green) on URD-inferred transcriptional trajectory of intestinal epithelial cells (Sur et al 2023). (**B**) Notch signaling activity: Immunofluorescence against Tg(*Tp1-bglob:eGFP*) with HCR marking *best4+* cells (*best4*), anterior enterochromaffin cells (*penka*), and goblet cells (*muc2.1*). Dotted circles highlight TP1– *best4*+ cells. (**C–D**) Notch inhibition: Notch signaling (GFP immunofluorescence detecting *Tp1:eGFP*), *best4*+ cells (*best4* mRNA), enterochromaffin cells (*adcyap1a* mRNA), all enteroendocrine cells (GFP immunofluorescence detecting *neurod1:eGFP*), goblet cells (*muc2.1* mRNA), and tuft-like cells (*pou2f3* mRNA) in DMSO vehicle-treated or DAPT-treated (72–120 hpf) larvae. White arrowheads indicate TgBAC(*neurod1:eGFP*)+ enterochromaffin cells and yellow arrowheads indicate *pou2f3*^+^ tuft-like cells. (**E–G**) Notch signaling increase: HCR for *best4+* cells (*best4*), enterochromaffin cells (“EChR”, *adcyap1a*), and goblet cells (*muc2.1*) after heat-shocking Tg(*hsp:Gal4*); Tg(*UAS:myc-Notch1a-ICD*) or Gal4– siblings from 72–96 hpf. (**H–J**) *dll4* knockdown: HCR for *best4+* cells (*best4*), enterochromaffin cells (*adcyap1a*), and goblet cells (*muc2.1*) after injection of control morpholino or *dll4* morpholino. See also Figures S6 and S7.

We next tested whether Notch signaling is required for *best4*+ cell specification by treating larval zebrafish with DAPT, a γ–secretase inhibitor that prevents Notch receptor activation^64^. We applied DAPT at three distinct time windows – (i) during the absorptive versus secretory decision (32–45 hpf)^29,31,32^, (ii) when the intestinal bulb expands and the first secretory cell types arise (48–72 hpf)^31,32^, and (iii) after secretory progenitors are specified, when *best4*+ cells are emerging (72–120 hpf)^16^ (Fig. 4C–D, Fig. S6D–N). Notch inhibition was confirmed by reduced GFP expression in the Tg(*Tp1-bglob:eGFP*)^um14^ Notch reporter line (Fig. 4C). Consistent with Notch’s role in suppressing secretory progenitor specification, Notch inhibition from 32–45 hpf increased the number of all secretory derivatives, including goblet cells (*muc2.1*^+^), enterochromaffin cells (*adcyap1a*^+^/*penka*^+^), *best4*+ cells, and tuft-like cells (*pou2f3*^+^) (Fig. S6D–F, J, N), though this increase was not statistically significant for *best4*+ cells (Fig. S6J, N). Inhibition of Notch signaling from 48–72 hpf similarly led to a modest, non-significant increase in all secretory derivatives (Fig. S6G–J, N). Importantly, DAPT treatment from 72–120 hpf only affected some secretory derivatives. First, we observed that it led to a strong reduction in *best4*+ cells, consistent with the hypothesis that Notch signaling is important for *best4*+ cell specification (Fig. 4C–D). Second, it led to a corresponding increase in the number of enteroendocrine cells (EECs), marked by TgBAC(*neurod1:GFP*)^nl1^ (Fig. 4C–D). Goblet and tuft-like cell numbers were not significantly altered (Fig. 4C–D). Since EECs are a heterogeneous population, we investigated whether a particular EEC subpopulation was increased during Notch inhibition. First, staining for *adcyap1a*, which marks enterochromaffin cells (the largest EEC subpopulation) revealed that the overall increase in EECs is dominated by the increase in enterochromaffin cells (Fig. 4D). Second, staining for *penka,* which marks an anterior subset of *adcyap1a*^+^ enterochromaffin cells showed an increase and distal extension of *penka*^+^ enterochromaffin cells when Notch is inhibited (Fig. S6K–N).

We then tested whether Notch signaling is sufficient to drive *best4*+ cell specification by upregulating Notch signaling via heat-shocking Tg*(hsp:Gal4);* Tg*(UAS:myc-Notch1a-ICD)* larvae when *best4*+ cell specification is ongoing (Fig. 4E–G). Notch upregulation between 72–96 hpf increased *best4*+ cells and reduced *adcyap1a*^+^ and *penka*^+^ enterochromaffin cells (Fig. 4E–G, Fig. S6O–R), which is the inverse effect of Notch inhibition. As expected, goblet cell numbers were not significantly altered by increased Notch signaling (Fig. 4E–G, Fig. S6O–R).

Altogether, these results demonstrate that *best4*+ cells receive Notch signaling at the time of their specification, require Notch signaling for specification, and can be increased by upregulating Notch signaling, demonstrating that Notch is a key developmental signal that governs specification of *best4*+ cells.

### *dll4* is a key Notch ligand involved in *best4*+ cell specification

Notch receptors are typically stimulated by ligands from the Delta or Jagged families^65–67^. To determine which ligands are expressed in intestinal populations when *best4*+ cells are specified, we used our previous scRNAseq dataset^16^. We observed strong expression of *dll4* and *dld/DLL1*, low levels of expression of *dla/DLL1*, *dlb/DLL3*, and *dlc/DLL3*, and no expression of Jagged family members (Fig. S7A). *dll4* was the most highly expressed Notch ligand in intestinal cells, with low expression levels in *best4*+ cells and high levels in enterochromaffin cells (Fig. S7A). Since we observed that Notch affected *best4*+ cells and enterochromaffin cells oppositely (Fig. 4D, G, Fig. S6N, R), we focused our experiments on *dll4*. Since Delta only signals to neighboring cells, we first performed HCR for *dll4* and *best4* alongside the Tg(*Tp1-bglob:eGFP*)^um14^ Notch reporter to determine whether *dll4+* cells are physically proximal to *best4*+ cells during their specification. We observed Tp1:GFP+ cells with varying levels of *best4*+ expression (*i.e.* cells actively receiving Notch signaling in the process of activating *best4* expression) adjacent to *dll4*+ cells, which is consistent with a possible role for Dll4 in the Notch signaling that specifies *best4*+ cells (Fig. S7B, arrowheads). To test the role of *dll4*, we then knocked down *dll4* expression using two splice-blocking *dll4* MOs. Both *dll4* MOs resulted in a significant reduction of *best4*+ cells and a concurrent increase in the number of enterochromaffin cells (*adcyap1a*^+^) (Fig. 4H–J, Fig. S7C–H), recapitulating the DAPT-mediated Notch inhibition phenotype (Fig. 4D, G, Fig. S6N, R). Interestingly, we also observed a reduction in goblet cells (Fig. 4H–J, Fig. S7C–H) which were unaffected during Notch receptor manipulations (Fig. 4D, G, Fig. S6N, R). This effect was modest when goblet cells were identified via an early marker^33^ (*agr2,* Fig. 4H–J) and more extreme when identified via a mature differentiation marker (*muc2.1*, Fig. S7C–H), suggesting effects on both specification and maturation, though the mechanism remains unclear. Altogether, these observations identify that *dll4* is a key ligand involved in *best4*+ cell specification, and that it is expressed lowly in some secretory progenitors and *best4*+ cells and strongly in EECs. During *best4*+ cell specification, *dll4* might be provided by mature enterochromaffin cells, by enterochromaffin cells born alongside *best4*+ cells, or even by *best4*+ cells themselves, as explored in the Discussion.

### *meis1b* confers *best4*+ cell identity

Next, we aimed to identify key elements of the gene regulatory network that confers *best4*+ cell identity within secretory progenitors that have received Notch signaling. Our previous developmental trajectory identified *meis1b* as one of the earliest *best4*+ cell-specific TFs (Fig. 5A, Fig. S8A), and *MEIS1* is also expressed in *best4*+ cells across species (Fig. 1B), suggesting it is highly important. To functionally test the role of *meis1b in vivo*, we created a *meis1b* whole-gene deletion using CRISPR/ Cas9 (Fig. 5B). We confirmed that *meis1b^y730^* is a null allele, since the gene is completely deleted, and there is no residual expression of *meis1b* mRNA in 6 dpf *meis1b*^−/−^ zebrafish larvae (Fig. S8B). *meis1b* mutants do not inflate their swim bladder, and interestingly, also have an inner ear phenotype (Fig. S8C, Sven Reischauer and Olov Andersson, personal communication), which we observed also in *meis1b^y730^* and used to sort *meis1b*^−/−^ larvae for downstream experiments.

**Figure 5:**
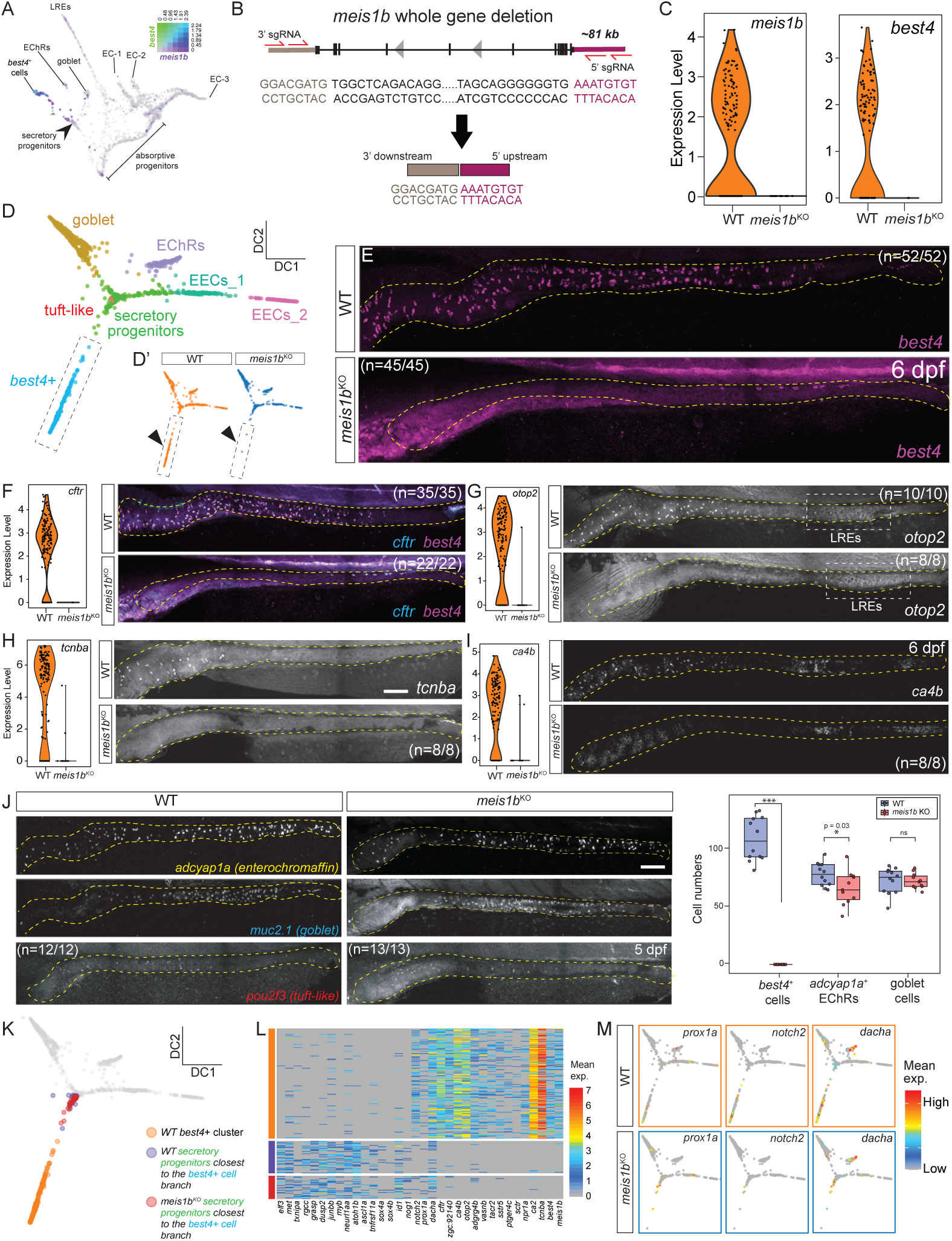
*meis1b* is essential for *best4*+ cell specification. (**A**) Co-expression of *meis1b* (purple) and *best4* (green) on URD-inferred transcriptional trajectory of intestinal epithelial cells (Sur et al 2023). (**B**) Nature of *meis1b* whole gene deletion. (**C**) Loss of *meis1b* and *best4* expression in secretory derivatives in *meis1b*^−/−^ mutant scRNAseq. (**D**) Diffusion map of all intestinal secretory derivatives colored by cluster (**D**) and genotype (**D’**). (**E–I**) Loss of expression of *best4*+ cell markers, including *best4* (E), *cftr* (F), *otop2* (G), *tcnba* (H), and *ca4b* (I) in *meis1b* mutants, shown via HCR and violin plots of expression within secretory derivatives. Scale bar: 100 um. (**J**) HCR of enterochromaffin (*adcyap1a*), goblet (*muc2.1*), and tuft-like cell (*pou2f3*) markers in *meis1b*^−/−^ mutants and WT siblings. Scale bar: 100 um. (**K**) Diffusion map showing wild-type *best4*+ cells (orange) and the secretory progenitors nearest *best4*+ cells (distance in diffusion map space) from wild-type (blue) and *meis1b* mutant (red) samples. (**L–M**) Differentially expressed genes (manually curated) between *meis1b* mutant secretory progenitors (blue) and either WT *best4*+ cells (orange) or WT secretory progenitors (red), shown via heatmap (L) or on diffusion map (M). See also Figures S8 and S9.

To investigate the role of *meis1b* in *best4*+ cell development, we performed scRNAseq on dissected intestines from 6 dpf *meis1b^y730^* mutant zebrafish larvae and their wild-type (WT) siblings (Fig. S8D–E). *meis1b^y730^* mutant transcriptomes indeed lacked *meis1b* expression and had not upregulated the paralogous gene *meis1a* (Fig. 5C, S8F). *best4* expression was eliminated entirely (Fig. 5C), and no *meis1b* mutant cells clustered with wild-type *best4*+ cells (Fig. 5D–D’, Fig. S8D–E). We confirmed via HCR that *best4* expression was eliminated in the intestine (Fig. 5E). Additionally, expression of other characteristic *best4*+ cell markers was also eliminated in both scRNAseq and HCR assays, including *cftr*, *otop2*, *tcnba*, and *ca4b* (Fig. 5F–I). The loss of numerous *best4*+ cell markers shows that *meis1b* is required for *best4*+ cell development. Goblet cells and tuft-like cells did not change in *meis1b*^−/−^mutants, and enterochromaffin cells were decreased only modestly (Fig. 5J). This suggests that secretory progenitors are not perturbed in *meis1b^y730^* mutants, since other secretory derivatives formed properly. Moreover, the effect of *meis1b* loss of function does not phenocopy Notch inhibition (in that *best4*+ cells are affected without an opposing effect on enterochromaffin cells), suggesting that *meis1b* acts downstream of Notch signaling.

We next identified which aspects of the *best4*+ cell developmental program were still expressed in *meis1b^y730^* mutants by identifying *meis1b* mutant secretory progenitor cells whose transcriptomes were biased towards wild-type *best4*+ cells, based on their proximity to wild-type *best4*+ cells in a diffusion map — a deterministic, nonlinear form of dimensionality reduction^68^ (Fig. 5K; Fig. S9A–B). *meis1b* mutant secretory progenitors were nearly transcriptionally identical to wild-type secretory progenitors, retaining expression of progenitor markers, including *ascl1a*, *atoh1b*, *tnfrsf11a, grasp, and dusp2* (Fig. 5L; Fig. S9C–E). *meis1b* mutant secretory progenitors also lacked expression of nearly every *best4*+ cell marker, with the exceptions of *prox1a*, *dacha*, and *notch2* (Fig. 5M). *notch2* was expressed at lower levels than typical for wild-type *best4*+ cells (Fig. 5L–M), suggesting *meis1b* may regulate *notch2* in a forward feedback loop. Since (1) *meis1b* mutants have unaffected secretory progenitors, (2) *meis1b* mutants do not phenocopy a loss of developmental signal, and (3) *meis1b* is expressed very early in *best4*+ cell development, we propose that *meis1b* is the critical factor that confers *best4*+ cell identity after Notch signaling, and is accordingly required for expression of nearly all *best4*+ cell markers.

### *pbx3a* is required for spatially heterogeneous gene expression in *best4*+ cells

We also previously identified *pbx3a* as an extremely specific *best4*+ cell marker at 5 dpf^16^ (Fig. 6A, Fig. S8A) and observed that Pbx-family members are expressed in *best4*+ cells in most animals (Fig. 1B). Our previous trajectory analysis suggested that *pbx3a* potentially turns on later than *meis1b*^16^ (Fig. 6A). To assess the functional role of *pbx3a in vivo*, we generated a *pbx3a* whole gene deletion mutant using CRISPR/Cas9 (Fig. 6B). Successful deletion of the *pbx3a* gene was confirmed using Sanger sequencing and loss of *pbx3a* mRNA signal in 6 dpf *pbx3a*^−/−^ larvae (Fig. 6B–D). In contrast to the complete loss of *best4*+ cells observed in *meis1b* mutants, *pbx3a^y731^* mutants retained *best4*+ cells (Fig. 6C–D), indicating that *pbx3a* is not required for *best4*+ cell specification.

**Figure 6:**
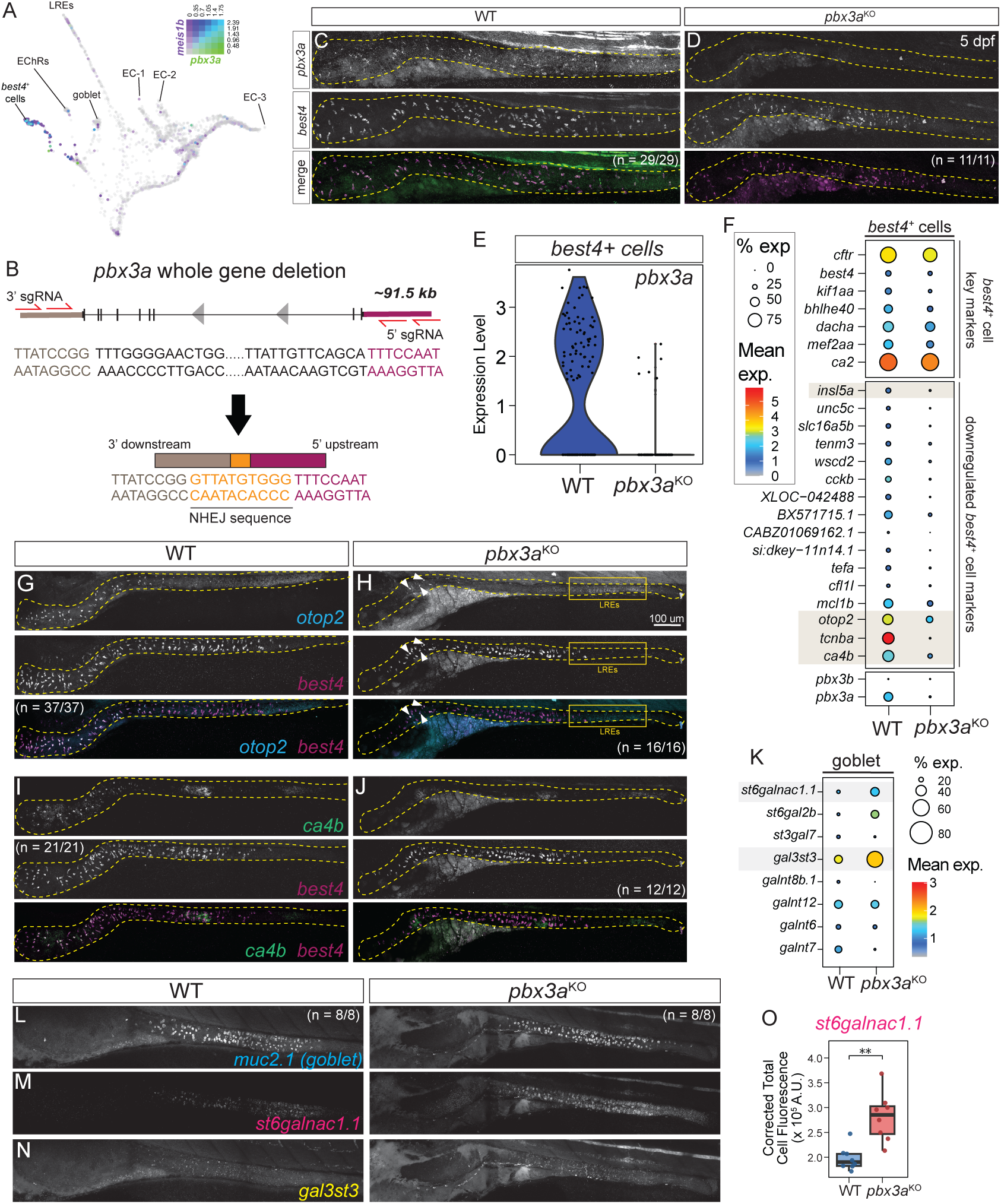
*pbx3a* is a regulator of regionalized *best4*+ cell identity. (**A**) Co-expression of *meis1b* (purple) and *pbx3a* (green) on URD-inferred transcriptional trajectory of IECs (Sur et al 2023). (**B**) Nature of *pbx3a* whole gene deletion. (**C, D**) HCR of *pbx3a* and *best4* in *pbx3a* mutants and WT siblings. (**E**) Reduced *pbx3a* expression in *pbx3a* mutant scRNAseq. (**F**) Top differentially expressed genes in *best4*+ cells between WT and *pbx3a* mutants. (**G–J**) HCR of selected *best4*+ cell genes differentially expressed in *pbx3a* mutants: *otop2* (**G, H**) and *ca4b* (**I, J**). Arrowheads indicate rare residual expression overlapping *best4*. (**K**) Top differentially expressed genes within goblet cells between WT and *pbx3a* mutants. (**L–O**) HCR of selected goblet cell genes differentially expressed in *pbx3a* mutants: *st6galnac1.1* (**M**) and *gal3st3* (**N**). (**O**) Corrected total cell fluorescence (y-axis) of *st6galnac1.1* HCR within a 100 x 100 pixel mid-intestinal region in *pbx3a* mutants and wild-type siblings. See also Figure S10.

To identify *pbx3a* targets in *best4*+ cells, we performed scRNAseq on dissected intestines from *pbx3a^y731^* mutant zebrafish larvae and their WT siblings (Fig. S10A, B). *pbx3a* mutants did not upregulate *pbx3b*, indicating a lack of genetic compensation in the *pbx3a^y731^* allele^69,70^, and had a transcriptionally distinct *best4*+ cell population, consistent with our observations from HCR (Fig. S10C, dotted circle). Only a few cells were *pbx3a*+, suggesting that there was very minor contamination of wild-type or heterozygous animals in our mutant sample (Fig. 6E, Fig. S10C). Nonetheless, differential expression analysis identified several *best4*+ cell-specific markers with altered expression in *pbx3a* mutants, including reduction of *otop2*, *ca4b*, *tcnba*, *wscd2,* and *insl5a* as well as increase of *satb2*, *adgrg6*, *cldn15a*, and *aldh9a1a.1* (Fig. 6F, Fig. S10D–F). Interestingly, most of these *pbx3a* targets are spatially restricted within the zebrafish intestine (Fig. 2J), though they do not have a consistent spatial expression pattern, and *pbx3a* seems to activate some while repressing others. Some spatially regionalized *best4+* cell markers are not regulated by *pbx3a*, since *ca2*, *dacha*, *cfd*, *vasnb*, and *tacr2* were unaltered in *pbx3a^y731^* mutants (Fig. 6F). To confirm our observations, we stained for some anteriorly restricted (e.g., *tcnba, otop2*, *ca4b*) and mid/posterior-restricted putative *pbx3a* targets (e.g., *insl5a*; Fig. 6G–J, Fig. S10G–H) in *pbx3a*^−/−^ mutant larvae. This demonstrated complete loss of *tcnba* in the gut, consistent with our scRNAseq analysis (Fig. S10E, G). For genes expressed in *best4*+ cells and another cell type, *pbx3a* only affected their expression in *best4*+ cells: *otop2* was nearly eliminated from *best4*+ cells but remained expressed in lysosome-rich enterocytes (Fig. 6G–H), *ca4b* was eliminated from *best4*+ cells although retained its low-level expression in enterocytes (Fig. 6I–J), and *insl5a* was eliminated from *best4*+ cells but remained expressed in EECs (Fig. S10H). Interestingly, in *pbx3a* mutants, a few rogue *best4* cells continued to express *otop2* (Fig. 6H, arrowheads). These results suggest that *pbx3a* is not required for *best4*+ cell specification but instead shapes spatially defined gene expression patterns within *best4*+ cells, acting in the anterior to activate several anterior markers and repress several putatively posterior markers, though some of these regulatory relationships may be indirect. The infrequent persistence of *otop2* expression in *pbx3a^y731^* mutants, the multiple distinct expression patterns that require *pbx3a,* and its potential dual involvement in activation and repression implies that *pbx3a* may not provide spatial information itself, but instead may work to potentiate the action of additional spatially restricted factors.

The gut functions as a complex, integrated ecosystem where various cell types are in constant and dynamic crosstalk to maintain homeostasis. Thus, we also performed differential expression analysis to identify whether *pbx3a* mutants exhibited non-autonomous transcriptional changes in IEC populations that do not express *pbx3a*. While most cell types were unaffected, goblet cells had altered gene expression in *pbx3a^y731^* mutants (Fig. 6K). *muc2.1* expression was modestly elevated in *pbx3a*^−/−^larvae, but we observed stronger upregulation of enzymes involved in post-translational mucin modification, including *gal3st3* (an O-glycan sulfotransferase) and *st6galnac1.1* (a sialyltransferase). Consistent with our transcriptomic data, HCR analysis confirmed that goblet cell numbers were unaffected in *pbx3a*^−/−^ mutants, but they exhibited elevated expression of *gal3st3* and *st6galnac1.1* (Fig. 6K–O). Since *best4*+ cells are the only IECs that express *pbx3a*, and it is not expressed in neighboring tissues (e.g. intestinal smooth muscle or the immune system), these findings raise the exciting possibility that *best4*+ cells influence the behavior of other cell types in the intestine, though the mechanisms and functional effects remain to be discovered.

### *best4*+ cells do not absorb fat or proteins

Despite our findings that *best4*+ cells descend from the secretory cell lineage, they express several markers of absorptive cells (and were in fact initially described as enterocytes^1,3,17,28,35,35,36^. Thus, we tested whether *best4*+ cells have any absorptive activity. To test whether *best4*+ cells absorb protein, we microgavaged 7 dpf Tg(*best4:eGFP, cryaa:mCherry*)^y7^^27^ larvae with the fluorescent protein, mCherry (Fig. 7A). 1-hour post-gavage, no mCherry fluorescence was observed in GFP+ cells (Fig. 7B–C), though significant absorption of mCherry protein had occurred in lysosome-rich enterocytes (LREs) which are specialized for protein absorption. To test whether *best4*+ cells absorb long-chain fatty acids, we microgavaged 7 dpf Tg(*best4:mApple, cryaa:eGFP*)^y726^ larvae with fluorescent lipids (BODIPY-FL C16). 1-hour post-gavage, several cells were marked by BODIPY-FL C16 fluorescence, all in the anterior bulb, though none were mApple+ (Fig. 7D–E, yellow arrowheads). Though several potential nutrients remain to be tested, altogether, these results suggest that *best4*+ cells are not involved in nutrient uptake.

**Figure 7:**
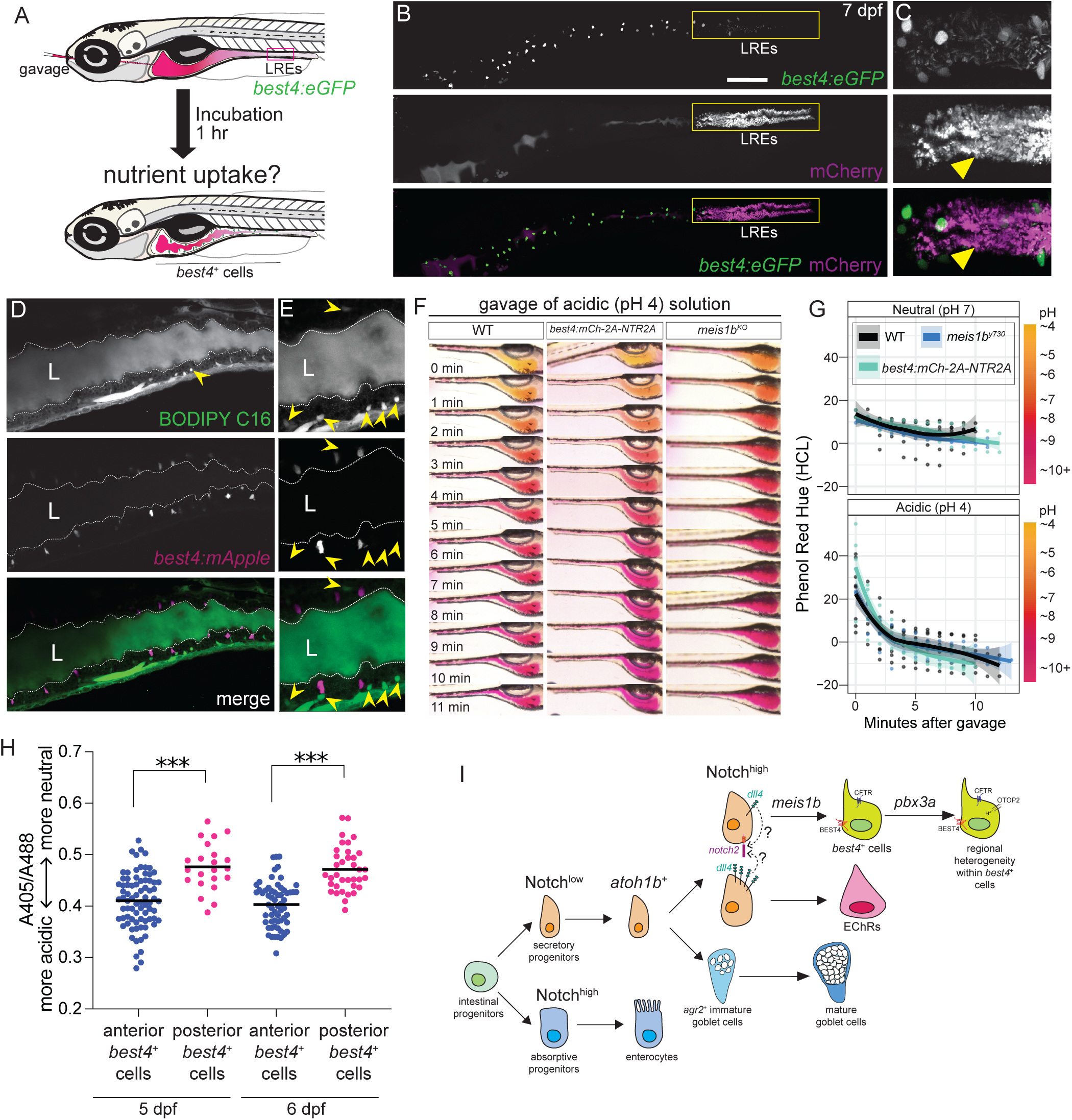
*best4*+ cells do not absorb nutrients, but have heterogeneous intracellular pH. (**A**) Microgavage technique for studying nutrient uptake in zebrafish intestines. (**B–E**) Live confocal images of a 7 dpf larval zebrafish intestine following gavage and internalization of mCherry protein (**B–C**) and BODIPY C16 (**D–E**). Yellow boxed regions: LREs. Yellow arrowheads: cells with mCherry or BODIPY C16 uptake. (**F–G**) Images and color quantification after microgavage of neutral (pH 7) or acidic (pH 4) 1% phenol red into wild-type, 0.1 mM metronidazole treated Tg(*best4:mCh-2A-NTR2A*), or meis1b^−/−^ mutant larvae. (**H**) 405/488 absorbance ratio of cytoplasmic fluorescence in Tg(*best4:pHluorin2*) 5–6 dpf larvae. Each point represents a cell. N = 6. (**I**) A proposed model of zebrafish *best4*^+^ cell specification and specialization. See also Figure S11.

### *best4*+ cells are not required to restore gut pH after acidic challenge

Since *best4*+ cells have only been recently molecularly described, their function and the *in vivo* consequences of their depletion remain a mystery. Our efforts to understand the development of these cells provided us with the genetic tools to assay changes in the intestine after preventing the birth of *best4+* cells using *meis1b^y730^* mutants or after eliminating *best4*+ cells by genetically ablating them using the *best4* enhancer. To ablate *best4*+ cells, we made Tg(*best4:mCherry–2A–NTR2A, cryaa:eGFP*)^y728^ zebrafish, which express a modified nitroreductase enzyme that converts metronidazole from a non-toxic precursor into a highly toxic drug specifically within *best4*+ cells, triggering their apoptosis^71–73^ (Fig. S11A). We used these tools to test previously proposed functions for *best4*+ cells.

Since *best4*+ cells express several ion channels, we hypothesized that they may contribute to pH regulation of the intestinal lumen. To test this, we microgavaged zebrafish larvae with pH indicator solutions (phenol red or m-cresol purple) that were initially either neutral or acidic and then imaged the color of the intestinal lumen over time as a measure of luminal pH. After introduction of either neutral (pH 7) or acidic (pH 4) phenol red solutions, color quickly shifted to magenta within 10 minutes, suggesting zebrafish larval intestines maintain a basic pH as previously noted^39^ and actively restore pH homeostasis after challenge (Fig. 7F–G, Fig. S11B). Similar results were obtained after introduction of acidic m-cresol purple, another pH indicator (Fig. S11C). The homeostatic pH of ZF larvae intestines is slightly alkaline (Fig. 7F–G), as previously noted^39^, and consistent with zebrafish’s lack of an acidified stomach^74,75^. Removal of *best4*+ cells, either through ablation using Tg(*best4:mCherry-2A-NTR2A*) or developmental perturbation (*meis1b^y730^* mutants) did not affect the intestine’s ability to return to homeostatic conditions or the time needed to do so (Fig. 7F–G, S11C–D). This suggests that either *best4*+ cells do not regulate luminal pH, or that multiple cell types in the intestine can perform this function, such that *best4*+ cells are not required.

### *best4+* cells exhibit heterogeneity in intracellular pH *in vivo*

Though zebrafish intestines can respond to acidic challenge in the absence of *best4*+ cells, it remained possible that *best4*+ cells respond to pH, rather than influence it. Moreover, previous *in vitro* experiments suggested that *best4*+ cells may acidify their intracellular pH in an acidic external environment^2,14^. Thus, we generated Tg(*best4:pHluorin2, cryaa:eGFP*)^y729^ zebrafish, which express a pH-responsive fluorescent protein^49–51^ in the cytoplasm of *best4*+ cells. *otop2* is an ion channel that imports protons, can alter intracellular pH^76,77^, and is only expressed in anterior *best4+* cells. Thus, we hypothesized that *in vivo*, anterior *best4*+/*otop2*+ cells might be capable of intracellular acidification and therefore exhibit a lower intracellular pH than posterior *best4*+ cells. To measure intracellular pH, we computed the 405/488nm fluorescence ratio of pHluorin2-positive *best4*+ cells located in the anterior bulb and posterior intestine (Fig. 7H). Anterior *best4*+ cells typically exhibited a more acidic intracellular pH than posterior *best4*+ cells. This suggests that *in vivo*, *best4*+ cell intracellular pH is spatially heterogenous, potentially shaped by the surrounding environment.

## DISCUSSION

### A model for zebrafish intestinal *best4*+ cell development

In this study, we test the developmental signals that specify *best4*+ cells and provide the first *in vivo* functional tests to determine the progenitors of *best4*+ cells and the TFs that regulate their development. Because these experiments were conducted *in vivo*, they bypass the need for artificially supplied developmental cues characteristic of human organoid systems and provide a complementary, in-context view of *best4*+ cell biology. In aggregate, our results identify the first comprehensive developmental framework and the beginnings of a gene regulatory network for the enigmatic *best4*+ cells. We propose that the following sequence of events specifies *best4*+ cells during intestinal development (Fig. 7I). First, Notch signaling regulates whether early intestinal progenitors adopt absorptive or secretory fates. Initially low Notch signaling specifies secretory progenitors^29,30,32^ (Fig. 3N-Q), some or all of which express *atoh1b*, similar to other animals^61,66,78^. Our study shows that *atoh1b*+ secretory progenitors give rise to larval *best4*+ cells (Fig. 3C–F) after Notch signaling acts a second time (most likely through *notch2*) to specify them (Fig. 4). Enterochromaffin cells also descend from *atoh1b*+ progenitors (Fig. 3I), and when Notch signaling is increased or decreased (either through its receptor or ligand), enterochromaffin cell numbers are altered in the opposing direction to *best4*+ cells (Fig. 4D, G, H), suggesting a terminal decision between these fates. Notch engagement by the ligand Dll4 is required for *best4*+ cell specification (Fig. 4H–J). While the key source of Dll4 remains untested, *dll4* is expressed strongly within EECs and weakly in *best4*+ cells themselves. One parsimonious model is that sister cells from a secretory progenitor division undergo lateral inhibition, where one upregulates Notch and the other upregulates Dll4, driving them to separate *best4*+ and enterochromaffin cell fates, though other potential models remain (as discussed below). Notch signaling within secretory progenitors activates the downstream TF *meis1b,* which confers *best4*+ cell identity and directly or indirectly inhibits secretory progenitor transcription factors (e.g. *atoh1b*) which decline during *best4*+ cell differentiation, but not in *meis1b* mutants (Fig. 5). Specified *best4*+ cells then respond to additional spatial cues that cooperate with *pbx3a* to establish regional heterogeneity within the lineage (Fig. 6). Regionalized gene expression may then establish functional heterogeneity, such as differences in intracellular pH (Fig. 7H).

This model introduces several additional developmental questions. First, the transcriptional heterogeneity of secretory progenitors is not well defined in zebrafish or mammals, and it is unclear whether secretory progenitor TFs act hierarchically to sequentially exclude particular secretory derivatives or merely act combinatorially to bias equipotent progenitors toward different fates. Recent results in mice suggest that not all secretory progenitors express Atoh1^79^, and recent studies in rats have shown that Prox1 is expressed in developing *best4*+ cells^16^. While the role of *prox1* remains to be tested definitively, its conserved expression in *best4*+ cells raises the possibility that *atoh1b* and *prox1a* co-expression may define the progenitors competent to become *best4*+ cells. This would be consistent with our observation that zebrafish *best4*+ cells and EECs are the two cell types with *atoh1b* and *prox1a* in common^16^ (Fig. 3G, I; Fig. S8A), and Notch signaling coordinately affects their numbers. This scenario, however, would necessitate investigation into the regulation of *atoh1b* and *prox1a* expression and whether they act sequentially or in parallel.

Moreover, while we propose potential Notch-mediated lateral inhibition between *best4*+ and enterochromaffin cells, Notch signaling might act via other mechanisms. For instance, studies in the *Xenopus* mucociliary epithelium propose that graded, dose-dependent Notch signaling specifies multiple cell fates simultaneously while global Notch signaling regulators tune regional cell type proportions^80,81^. Additionally, recent studies in mouse intestinal organoids suggest that different frequency oscillations of Notch activation specify different intestinal fates. Prior studies of zebrafish secretory progenitor specification identified two distinct temporal windows of Notch activity^82^, which could implicate this mechanism in zebrafish development as well. Finally, low levels of *dll4* expression in *best4*+ cells could trigger Dll4–Notch2 *cis*-interactions which can counter-intuitively strengthen Notch signaling^83,84^, raising the possibility that autocrine amplification of Notch activity through Notch2 may be crucial for *best4*+ cell fate. More sophisticated manipulations and assays will be needed in the future to test these additional models, but potentially one could explain an experimental discrepancy that remains a mystery to us—namely that *dll4* knockdown reduces goblet cell numbers, but global Notch inhibition seemingly does not.

### Conservation of developmental regulation

The signaling pathways (*notch2*) and TFs (*meis1b*, *pbx3a*) that we implicate in *best4*+ cell development in this study are conserved across most or all vertebrate species (Fig. 1B), thus we predict that they are broadly important. High levels of *notch2* expression have even been found in *best4*+ cells of sturgeons, an ancient pre-Jurassic teleost lineage often considered a living fossil^19^. Our work agrees with previous studies in rats and human organoids that also found Notch signaling is essential for *best4*+ cell development^14,36^. We additionally show in an *in vivo* system that *best4*+ cells are exclusively responding to Notch at the time of their specification (Fig. 4B) and that increased Notch signaling is sufficient to trigger increased production of *best4*+ cells (Fig. 4E–G). However, the Delta ligands that specify *best4*+ cells might not be as conserved. We identify *dll4* as a key ligand involved in *best4*+ cell development, but recent human organoid studies identified DLL1 as a key ligand involved in colonic *best4*+ cell development^85^. Human organoid studies and transcriptional data suggest differences in the developmental programs of small intestinal and colonic *best4*+ cells; this ligand difference may reflect differences between species, or potentially that zebrafish larval *best4*+ cells develop more similarly to human small intestinal *best4*+ cells whose key ligands have not yet been determined.

Other studies have identified inputs into *best4*+ cell development from hematopoietic and immune-related factors. For instance, human colonic intestinal epithelium mutant for *SPIB* (a TF typically associated with hematopoiesis) lacks *best4*+ cells, M-cells, and tuft cells, though *SPIB* expression is less conserved than *meis1* expression in *best4*+ cells (Fig. 1B). Additionally, *best4*+ cell numbers change upon perturbation of the RANK/RANKL pathway in humans and zebrafish after challenge with IFNγ or *Vibrio*^36,86^, all of which suggest potential involvement of immune responses in determining overall *best4*+ cell numbers. It remains unclear whether these perturbations directly influence developmental regulation underlying *best4*+ cell specification, alter the makeup of secretory progenitors (thereby affecting the specification of multiple secretory derivatives simultaneously), or affect the behavior of another cell type that indirectly influences *best4*+ cell number. Work remains to identify how these immune derived cues might integrate with and influence the developmental pathway described in this study.

### Conservation of developmental origins

In this study, we provide the first direct *in vivo* lineage tracing evidence in any organism to determine the progenitors from which *best4*+ cells descend, validating our previous computational predictions^16^. We find that *best4*+ cells in all segments of the larval zebrafish gut arise from *atoh1b*^+^ secretory progenitors (Fig. 3B–F), and that *best4*+ cell numbers are significantly reduced after knockdown of *atoh1b* (Fig. 3J–M). Our findings align with previous work in zebrafish that identified a population of Notch-responsive secretory cells (NRSCs) that arise during larval development^87^, which are likely to be *best4*+ cells. Additionally, this work aligns with predictions from trajectory analyses within the human and rat small intestine^4,14^. Interestingly, developmental trajectories predict that human colonic *best4*+ cells have an absorptive origin, which is consistent with colonic organoid experiments that showed *best4*+ cells still form after *ATOH1* KO^41^. We cannot exclude the possibility that some *best4*+ cells have a different origin in larval zebrafish, since our lineage tracing approach does not label cells in perpetuity, but *best4*+ cells arising from different progenitors would not be confined to a specific intestinal region. The discrepancy between observed secretory and absorptive origins may (1) resolve as additional mammalian colonic experiments are performed, (2) reflect a difference in species, where zebrafish use an ancient pathway for *best4*+ cell formation that has been further expanded in humans, or (3) reflect a difference in developmental stage, since this study was conducted in larvae, whose intestines will further develop during juvenile stages.

### *best4*+ cell function and morphology

In addition to developmental signals and TFs, several characteristic functional genes are conserved among vertebrate *best4*+ cells, but *best4*+ cell functions remain largely untested *in vivo*, since functional assays and *in vivo* ablation systems for these cells remain limited. The conserved transcriptional program of *best4*+ cells suggests fluid and ion homeostasis as one of their primary functions. While mice lack *best4*+ cells, many characteristic *best4*+ cell functional genes are redistributed to other intestinal cell types (Fig. 1D), which suggests that a key purpose of *best4*+ cells may be to compartmentalize and thereby confer regulation to these functions. That regulation might be accomplished by signaling through the numerous hormone receptors expressed by *best4*+ cells, through the immune system (given findings from *RANK* and *SPIB* mutants)^17,36^, or by luminal pH or ion content itself through *best4*+ cell-specific ion channels that are not redistributed to other IECs in mice (*best4* and *otop2)* (Fig. 1D). We present the first *in vivo* genetic removal or ablation of these cells (Fig. 5, Fig. S11A), which revealed that in zebrafish, *best4*+ cells are not required to restore intestinal pH after acidic challenge (Fig. 7F–G) but have spatial heterogeneity in their intracellular pH that may reflect functional heterogeneity (Fig. 7H). This raises exciting questions about how *best4*+ cells respond to changes in the intestinal environment, including its pH, that can be further pursued using these ablation models.

In zebrafish, we observed that *best4*+ cells extend dynamic protrusions toward the lumen (similar to EECs), suggesting they sample environmental conditions, and also basolaterally towards either neighboring epithelial cells, the intestinal stroma, or the enteric nervous system, suggesting potential interactions with surrounding cell types (Fig. 2A–F). Indeed, *best4*+ cells are rich with hormone receptors and secreted hormones. Additionally, our study identifies potential non-autonomous effects on goblet cell gene expression that may alter mucus properties and thereby the microbiota or barrier function (Fig. 6K–O). Recent work in human organoids also suggests potential communication between *best4*+ cells and goblet cells^85^, and projections from *best4*+ cells have also been seen in other animals^13,36^. Finally, NRSCs (likely *best4*+ cells) have been proposed to engage in IGF/EGF signaling that influences proliferation rates within intestinal cells^87^, which aligns with *in vitro* studies that propose tumor-suppressing roles for the *best4* and *otop2* genes^26,27,88,89^. Future work remains to identify the signaling partners and signaling networks *best4*+ cells participate in, which signals are potentially mediated by the dynamic projections of *best4*+ cells, and how *best4*+ cells contribute to the larger network of intestinal functions that go awry in disease.

In conclusion, this study shows that many aspects of *best4*+ cell development and function might be conserved between zebrafish and other animals, emphasizing that studying *best4+* cells across many systems with different advantages will collectively advance our understanding of their biology. Given their broad conservation, they are likely to have an essential role that has been selected for across evolution, though their functions remain to be determined. Here, we present a comprehensive description of *best4*+ cell development, including identifying key elements of its gene regulatory network that are likely conserved across vertebrates. Understanding *best4*+ cell development will shed light on whether changes in *best4*^+^ cells during disease are a result of intestinal dysfunction or contribute to disease progression, as well as how *best4*+ cells might be therapeutically manipulated. Additionally, we demonstrate a model system where *best4*+ cells can be genetically manipulated, removed at will, and observed in real time in the systemic context of an entire animal, including an intestinal stroma, enteric nervous system, active immune system, and even microbiota. This lays the groundwork to address key questions about *best4*+ cell function in an organismal context.

### Limitations of the study

Our study characterizes *best4*+ cell development in the larval zebrafish intestine—a developmental window in which the gut is forming for the first time and all epithelial lineages are being specified *de novo*. Although by 5 dpf, intestinal folds have formed in the zebrafish gut and specialized cell types are already present, homeostatic renewal of IECs has not yet begun, and additional maturation, looping, and potentially further regionalization of the intestine will occur in the following 2–3 weeks. Further investigation will be needed to determine whether the same developmental program is reused in adult stages for homeostatic replacement of *best4*+ cells.

## Resource availability

Further information and requests for resources and reagents should be directed to and will be fulfilled by Jeffrey A. Farrell. Plasmids and transgenic zebrafish lines generated in this study will be shared by the lead contact upon reasonable request. In cases where international transport of the lines is too complicated due to import restrictions or other considerations, the Tol2 construct needed to recreate the transgenic line may be shared instead. Data is currently being uploaded to standardized repositories to be released upon publication, but is currently available by request.

## Acknowledgments

Funding was provided by the NICHD Intramural Program to J.A.F. (ZIAHD008997) and the K99/R00 Pathway to Independence Award to A.S. (1K99HD115786). Olov Andersson and Sven Reischauer for personally communicating the *meis1b* phenotype; Eleanor Zagoren and Kaelyn Sumigray for sharing in-press scRNAseq data; Dan Castranova for assistance with live imaging and zebrafish lines; Ajay Chitnis, Katie Drerup, Katherine Rogers, and Brant Weinstein for reagents and zebrafish lines; Edan Foley, Dan Goldman, Tatjana Piotrowski, and Lihua Ye for generous donation of reagents that did not end up presented in this manuscript; and James Gagnon, Alan Hinnebusch, Micaela Murphy, Katherine Rogers, and members of the Farrell Lab for critical feedback on this manuscript. This work utilized computational resources of the NIH HPC Biowulf cluster.

## Author Contributions

J.A.F and A.S. conceived the study. A.S. and J.A.F wrote the manuscript. A.S. made zebrafish transgenic lines and mutants. A.S., E.S., and M.K.P performed fish husbandry and genotyping. A.S., E.X.S, M.P.N, J.A.F, and Y.W performed data analysis. J.W.S and B.F. generated the *atoh1b* endogenous knock-in line. L.E.D. conducted electron microscopy. L.F. and Y.S. generated Xenopus scRNAseq data. A.S., M.P.N, M.K.P, and J.A.F performed intestinal dissections. A.S., M.P.N, and J.A.F collected single-cell transcriptomes and J.I. contributed to sequencing and Cell Ranger. E.X.S, A.S., and M.P.N performed other experiments, including RNA *in situ* hybridization. E.X.S. optimized microgavage and performed all gavage experiments.

## MATERIALS AND METHODS

### Experimental model and subject details

Zebrafish (*Danio rerio*) used in this study include: wild-type TL/AB (Tupfel long fin/AB hybrids, generated from separately maintained TL and AB breeding stocks), and several previously established transgenic lines as well as lines generated in this study on a TLAB background. Previously established transgenic lines used in this study include: Tg(*hsp:Gal4*)^kca4+/−^, Tg(*UAS:myc-Notch1a-ICD*)*^kca3^* ^90^, Tg(*Tp1-bglob:eGFP*)^um14^ ^63^, and TgBAC(*neurod1:eGFP*)^nl1^ ^91,92^. Transgenic lines Tg(*best4:mApple, cryaa:eGFP*)^y726^, Tg(*best4:eGFP, cryaa:mCherry*)^y727^, Tg(*best4:pHluorin2, cryaa:eGFP*)^y729^, Tg(*best4:mCherry–2A–NTR2A, cryaa:eGFP*)^y728^, and an endogenous knock-in line *atoh1b*^y732^ (referred to in this study as *atoh1b^atoh1b–p2A–mGreenLantern^* or atoh1b^mGL^) was generated as part of this study. This study also led to the generation of developmental TF mutants such as *meis1b^y730^* and *pbx3a^y731^* whole gene deletion mutants.

This study includes the use of live zebrafish vertebrate embryos. Animals were handled according to National Institutes of Health (NIH) guidelines. All zebrafish work was performed at the *Eunice Kennedy Shriver* National Institute of Child Health and Human Development (NICHD) Shared Zebrafish Facility, under animal protocol 23-001. The NICHD Animal Care and Use program also maintains full AAALAC accreditation, is assured with OLAW (D16-00602) and is currently registered with the US Department of Agriculture (USDA). At the developmental stages studied in this work, zebrafish sex is not yet determined, hence, sex was not considered as a biological variable in this study.

Zebrafish embryos were raised in petri dishes containing E3 medium in the dark at 28°C for up to 7 days without feeding. Breeding adults were housed in a large zebrafish-dedicated recirculating aquaculture facility (4 separate 22000 L systems) in 1.8L tanks at standard (28°C) temperature on a 14-hr light/10-hr dark cycle. Fry were fed rotifers and adults were fed Gemma Micro 3000 (Skretting) 1–2 times per day. Water quality parameters were routinely measured, and appropriate measures were taken to maintain water quality stability. Larvae used for collection of single-cell transcriptomes or staining were never fed.

### Curation and pre-processing scRNAseq data across species

We curated intestinal single-cell atlases generated by other groups, including *Homo Sapiens*^3,4^ colon (31347 cells) and small intestine (12400 cells), *Bos Taurus*^6^ (88,013 cells), *Sus Scrofa* (Bama miniature pigs)^8^ (913 cells), *Rattus norvegicus*^14^ (7362 cells)*, Xenopus tropicalis* (unpublished; 18412 cells), larval^16^ (489,686 cells) and adult^28^ (28143) *Danio rerio,* and *Mus Musculus*^49^ (17512 cells). Except cow and pigs, cell annotation information was used as provided in the original dataset. For the cow and pig dataset, the counts matrices were downloaded and were used to create Seurat objects. We normalized the read counts of each cell using Seurat (v4.1.0)^93^, identified genes with the top 2000 highest standard deviations as highly variable genes, performed PCA analysis, and identified significant PCs. A Uniform Manifold Approximation and Projection (UMAP) was calculated using 50 nearest neighbors and the 30 most significant PCs. Leiden clustering was performed for these two datasets using the top 30 PCs and multiple resolutions. Final clustering resolution was chosen based on biological intuition, considering that individual clusters represented individual, well-known, conserved cell types based on their marker gene expression.

To maintain consistency in analysis across datasets, all Seurat datasets were converted to v4.1.0 for downstream analysis. To normalize clustering across datasets for downstream analysis, clustering was curated in the following ways across datasets: First, each dataset was cropped to just include intestinal epithelial cells (IECs). Second, among IECs, cell types that are not conserved across species were removed. For example, Paneth cells and LGR5+ stem cells from mammalian datasets were not considered, since no similar counterparts have been described yet in zebrafish. Third, clusters were renamed to match across species. Altogether, each dataset was cropped to include transit amplifying (TA) cells (mammals) or secretory progenitors (zebrafish/Xenopus), enterocytes, goblet cells, enteroendocrine cells, *best4*+ cells, and tuft cells.

### Identification of a conserved *best4*+ cell specific program

To identify a conserved program of *best4*+ *cells* across species, we first calculated *best4*+ *cell–enriched markers from each dataset.* Marker genes for *best4+ cells* were identified using Wilcoxon Rank Sum Tests using the command Seurat::FindMarkers(test.use = “wilcox”, logfc.threshold = 0.25), selecting markers at least 1.28-fold more strongly expressed in *best4*+ cells with a significance of *p* < 0.01. To refine these results, we applied an additional filtering step using a custom function that selects markers based on a minimum difference in the percentage of cells expressing the gene between *best4*⁺ cells and all other cells (pct.1 – pct.2 ≥ 0.1). For each cluster, genes passing these criteria were retained as high-confidence marker genes, and the resulting gene lists were used for downstream cross-species comparisons. Marker genes from each species were then converted to human orthologs using the HGNC Comparison of Orthology Prediction (genenames.org/tools/hcop/). For each species, when calculating conservation, all orthologs were maintained (*i.e.* to account for gene paralogs and gene duplication events); for instance, zebrafish *pbx3a* and *pbx3b* were both considered orthologs of human *PBX3* and considered downstream.

To identify which *best4+* cell-enriched genes were conserved across species, we identified the number of organisms for which any paralog of each gene was considered a “marker” as described above. Genes that marked *best4*+ cells in ≥6 of 8 considered datasets were considered “conserved”, while genes that appeared in only one species were considered “species-specific”. In addition, this process was repeated separately across the mammalian datasets and aquatic datasets individually to identify genes conserved within only mammalian *best4*+ cells and only *best4*+ cells of aquatic species. The genes shown in plots of conserved genes were further manually expanded to include some orthologs (PBX1 and NOTCH3) that are enriched in *best4*+ cells of several species and may be functionally redundant with more “conserved” genes. As a metric of how specific each conserved gene is to *best4*+ cells, first the average gene expression was calculated for each cluster in each data set. Then, the specificity (i.e. the percentage of mRNA of each gene present in *best4*+ cells) was calculated by dividing the mean expression of each gene within *best4*+ cells by the total mean expression of each gene across all intestinal epithelial cell types and multiplying the result by 100. These specificity values and average expression were plotted to visualize the strength and conservation of *best4*+ cell markers across species. Genes were manually ordered and plotted as a heatmap using ggplot2.

### Generation of transgenic zebrafish lines

Transgenic lines were established using the standard Tol2 protocol^94^. For the Tg*(best4:eGFP*, *cryaa:mCherry)*, Tg*(best4:mApple*,*cryaa:eGFP),* Tg*(best4:phluorin2*,*cryaa:eGFP), and* Tg*(best4:mCherry-2A-NTR2A, cryaa:eGFP)* transgenes, a 2.3 kb sequence ∼4.1 kb upstream of the transcriptional start site was PCR amplified off of genomic DNA using primers 5’–CTCTGGCTCGGTTGAGAAGG–3’ and 5’– TGGCCAGTGTCAGTTTCACA–3’ and then TOPO TA cloned into a pENTR Gateway vector, using the pENTR™5’-TOPO TA Cloning Kit (Invitrogen, Cat#K59120) to generate plasmid JFP713 (*p5E-best4-2.3*). JFP713 (*p5E-best4-2.3*) was then recombined using LR Clonase II Plus (Invitrogen Cat# 12538120) with: (a) pKD001 pME-minprom_EGFP (Addgene #195955)^95^, Tol2Kit #302 p3E-SV40polyA^94^ and pDEST-Tol2pA;*cryaa:mCherry* (Addgene #64023)^96^ to generate JFP717 *pTol2-best4-minprom-EGFP, cryaa:mCherry-SV40polyA*, (b) pCK068 pME-minprom-mApple^97^, Tol2Kit #302 p3E-SV40polyA^94^, and pDEST-Tol2pA,*cryaa:eGFP* (Addgene #64022)^96^ to generate JFP718 *pTol2-best4-minprom-mApple,cryaa:eGFP-SV40polyA*, (c) pME-pHluorin2 (Addgene #73794)^98^, Tol2Kit #302 p3E-SV40polyA^94^, and pDEST-Tol2pA,*cryaa:eGFP* (Addgene #64022)^96^ to generate JFP747 *pTol2-best4-minprom-pHluorin2,cryaa:eGFP-SV40polyA*, and (d) pME-mCherry-2A-NTR2A (a gift from the Weinstein Lab), Tol2Kit #302 p3E-SV40polyA^94^, and pDEST-Tol2pA,*cryaa:eGFP* (Addgene #64022)^96^ to generate JFP724 *pTol2-best4:mCherry-2A-NTR2A,cryaa:eGFP-SV40polyA*.

Embryos were injected at the one-cell stage with 40pg Tol2 transposase mRNA (produced from Tol2Kit #396 pCS2-FA-transposase) and 25–30 pg of *pTol2-best4-minprom-EGFP,cryaa:mCherry-SV40polyA*, *pTol2-best4-minprom-mApple,cryaa:eGFP-SV40polyA*, *pTol2-best4-minprom-pHluorin2,cryaa:eGFP-SV40polyA,* and *pTol2-best4:mCherry-2A-NTR2A,cryaa:eGFP-SV40polyA* plasmids. Injected F0 embryos from three independent clutches were assessed between 5–6 days post fertilization (dpf) and screened for *crystallin* (*cryaa*) expression in the zebrafish eye and mosaic fluorescent expression in the intestine. Gut fluorescence was observed under a Nikon Eclipse Ti2 inverted confocal microscope with a Nikon DS-R*i*2 camera. Injected mosaic animals were compared to uninjected control embryos, which did not exhibit any fluorescence in the eye or intestine, and were then transferred to the aquaculture facility. Upon reaching sexual maturity, F0 injected adults were outcrossed to TL/AB and their F1 progeny were screened for *cryaa*-driven expression in the eye and *best4*-driven expression in the intestine. F1 larval zebrafish exhibiting fluorescence in the eyes and gut were raised to adulthood and maintained as stable lines in the aquaculture facility. Expression of *best4:GFP* and *best4:mApple* in *best4*+ cells was confirmed using RNA *in situ* hybridization combined with immunofluorescence against GFP and Apple.

### Electron Microscopy

#### Sample preparation for Immuno-gold Electron Microscopy

Zebrafish intestines (7dpf; n = 50–70) were dissected from Tg(*best4:eGFP, cryaa:mCherry*)^y727^ zebrafish larvae and fixed with 4% paraformaldehyde (Electron Microscopy Sciences #157-8-5G) in phosphate buffer saline (PBS). Several fixed intestines were enrobed in 4% Agarose, and 100-micron Vibratome sections were produced. Sectioned intestines were rinsed in 50mM glycine in PBS for 25 minutes and then blocked with 2% BSA with 0.1% Saponin in PBS for 45 minutes. Next, sections were incubated for one hour at room temperature (RT) in Rabbit anti-GFP primary antibody (1:200, Rockland Cat# 600-401-215). Sections were then rinsed in incubation buffer (2% BSA with 0.05% Saponin in PBS) for 3 x 5 minutes each. Sections were then incubated in gold (1.4nm) Nanoprobes Alexa-Fluor 647 Fab anti-Rabbit secondary antibody (1:50) for 1 hour at RT. Incubated sections were rinsed in PBS, 3 x 5 minutes each. Vibratome sections were imaged using a Nikon Eclipse Ti2 inverted confocal microscope with a Nikon DS-R*i*2 camera to verify that the immunostaining enhanced the GFP signal. After imaging, the sections were post-fixed in 2% glutaraldehyde made in PBS for 30 minutes at RT on a rocker. Next, sections were rinsed in MilliQ water for 5 x 5 minutes each.

#### Silver Enhancement and processing for Electron Microscopy

Sections were transferred to Nanoprobes HQ silver enhancement solution and incubated at room temperature for 10 minutes. The enhancement was stopped by washing sections several times in MilliQ water. Sections were changed into 0.1M phosphate buffer (PB) and treated with 0.2% osmium tetroxide in PB for one hour on ice. Next, sections were rinsed in 0.1M PB on ice, 3 x 5 minutes each, followed by a rinse in 0.1M acetate buffer, pH 5.0 on ice, 2 x 5 minutes each. The sections were also en-blocked in 0.25% uranyl acetate made in 0.1M acetate buffer for one hour at 4°C. After rinsing the sections in 0.1M acetate buffer, pH 5.0, 3 x 5 minutes each the sections were processed using the variable wattage Pelco BioWave Pro microwave oven (Ted Pella, Inc., Redding, CA.) with the following steps: ethanol dehydration series up to 100% ethanol, followed by a Embed-812 epoxy resin (Electron Microscopy Sciences, Hatfield, PA.) infiltration series up to 100% resin. The epoxy resin was polymerized for 20 hours in an oven set at 60° C. Ultra-thin sections (90nm) were prepared on a Leica EM UC7 ultramicrotome. Ultra-thin sections were picked up and placed on 200-mesh copper grids (Electron Microscopy Sciences, Hatfield, PA) and post-stained with UranyLess (Electron Microscopy Sciences, Hatfield, PA) a uranyl acetate replacement and lead citrate. Imaging was performed on a JEOL-1400 Transmission Electron Microscope operating at 80kV and images were acquired on an AMT BioSprint 29 camera.

### Embryo pretreatment and fixation

Zebrafish larvae (4–7 dpf) were collected and fixed overnight in 4% paraformaldehyde at 4°C, then transferred to methanol and stored at –20°C. Prior to staining, larvae were rehydrated to PBST (phosphate buffered saline, 0.1% Tween-20 pH 7.3) through 3 graded methanol/PBST steps (75%:25% methanol:PBST, 50%:50%, 25%:75%, 100% PBST). After rehydration, samples were further permeabilized by proteinase K treatment (10 μg/mL in PBST) for 50–55-minutes, post-fixed in 4% paraformaldehyde for 20 mins at RT and then washed five times with PBST, for 5 mins each.

### Hybridization Chain Reaction (HCR) RNA in situ hybridization

HCR was performed for *best4^+^* cell specific markers as well as goblet cell and enterochromaffin cell markers. HCR probes were designed using the Özpolat lab probe generator^99^, available at <u>https://</u> <u>github.com/rwnull/insitu_probe_generator</u> following a similar workflow as described in our previous work^16^. For each gene, at least 20 probe pairs were ordered in OPools format (Integrated DNA Technologies) and resuspended in nuclease-free water to a working concentration of 1 µM. Oligos constituting probe sets used in this study are consolidated in Table S1. Following rehydration and fixation, subsequent steps were performed as previously described^16^.

### Immunostaining after HCR

Immunostaining was performed on *best4:eGFP* and *best4:mApple* to visualize *best4*+ cells and validate the corresponding transgenic lines, on *atoh1b^mGL^* larvae to identify cell types derived from *atoh1b^+^* progenitors, and on *Tp1:eGFP* to monitor Notch reporter activity. Following HCR, samples were permeabilized with 0.1% Triton X-100 in phosphate buffer saline (PBS) and subsequently blocked for 2 hours in blocking solution (5% Normal Donkey Serum (Jackson Labs), 10% Bovine Serum Albumin, 1% DMSO, and 0.1% Triton X-100 in phosphate buffered saline) at room temperature. Primary antibodies – anti-RFP (rabbit, Rockland 600-401-379) and anti-GFP (rabbit, Invitrogen A11122) were applied at 1:1000 in blocking buffer and rocked on a nutator overnight at 4°C. Following primary antibody exposure, samples were washed 6 times with PBST, 15 mins each Secondary staining was performed using goat Alexa Fluor 488-anti-rabbit (Molecular Probes, Cat#: A-31556) or goat Alexa Fluor 546-anti-rabbit (Molecular Probes, Cat#: A-11035) diluted 1:1000 in blocking buffer by gently rocking them overnight at 4°C. Secondary antibody incubation was followed by 6 washes in PBST, 15 mins each before preparing the samples for imaging.

### Microscopy and image analysis

Stained larvae were mounted in 1% low melting agarose diluted in 1X Danieau buffer (58 mM NaCl, 0.7 mM KCl, 0.4 mM MgSO_4_, 0.6 mM Ca(NO_3_)_2_, 5 mM HEPES, pH 7.6) and imaged on a Nikon Eclipse Ti2 inverted microscope with a Nikon DS-R*i*2 camera using a 40X long working distance water/ oil objective. Image processing was performed in ImageJ/Fiji^100^ and Photoshop (Adobe). The acquired z-stacks were projected in Fiji, and brightness and contrast adjustments were applied using linear LUT scaling in Fiji. Cropping and resizing was performed using Photoshop (Adobe) and final figures were assembled using Illustrator (Adobe). The elongated zebrafish gut often necessitated acquiring multiple fields of view with ∼10% overlap between each field of view, using the Nikon Elements Software tiling features. Post-processing to stitch them together was performed using the “File > Stitch large image from files” dialogue box with default parameters in the Nikon Elements Software that uses an Image Registration based precise stitching method.

### Generating an endogenous knock-in line

To generate the *atoh1b*^y732^ *(atoh1b^atoh1b–p2A-mGreenLantern^*) knock-in line, AB fish were sequenced to ensure homozygosity 40 bp upstream and downstream of the *atoh1b* stop codon, covering potential guide sequences and homology arms for a HDR template. An active crRNA (5’–atgtgtggtaatcagcgtcct–3’) that spanned the *atoh1b* stop codon was identified using fluorescent PCR to detect generation of indels^101^. An HDR template was generated by PCR amplification from a modified pJS178 dbh-3.0:CFP-P2A-Gal4ff plasmid, in which mGreenLantern had been subcloned downstream of the P2A sequence using the pKWR233 pCS2+-bmp2b-mGreenLantern plasmid (a gift from the laboratory of Katherine Rogers) as the mGreenLantern source, using 5′ AmC6 end-protected primers with sequence homology to the genomic region surrounding the *atoh1b* stop codon^102^: 5AmMC6-atttcagcgacactgaagaaggacacactggaggacgcTCCGGAGCCACGAACTTC (Integrated DNA Technologies) and 5AmMC6-ctataaatgactatagcatttaatgtgtggtaattacttgtacagctcgtccatgtcatg (Integrated DNA Technologies). For the knock-in injection, 1-cell stage embryos were injected into the cell with 24 fmol cRNA (Integrated DNA Technologies), 20 fmol Cpf1 (New England BioLabs), 343 mM KCl, and 98 picograms HDR template. Injected embryos were screened for hindbrain mGL expression at 24 hours post fertilization. Embryos showing strong expression were raised to adults and outcrossed to identify germline transmitters.

### Live Imaging of larval *best4*+ cells and *atoh1b*+ progenitors

Larval zebrafish were anesthetized with buffered tricaine and either mounted in 1% low-melt agarose (Aquapor LE) or placed in an imaging dish with a mold made of 2% agarose formed with a 3D printed stamp. Live confocal images were acquired using a Nikon Ti2 inverted microscope equipped with a Yokogawa CSU-W1 spinning disk confocal and a Hamamatsu Orca Flash 4 v3 camera (Video S3) as well as a Nikon DS-R*i*2 camera on a Nikon A1 microscope (Videos S1, S2). Images were captured every 15 mins on a 20X water immersion on the spinning disk confocal and on a 40X water immersion on the Nikon A1 confocal using Galvano settings.

### Morpholino injections

Previously published morpholinos (MO) targeting *dll4*^103,104^ were obtained from Gene Tools (Philomath, OR) and were resuspended in water. *atoh1b* MO was obtained as a gift from the Chitnis lab. To knock-down *dll4* and *atoh1b* function, two *dll4* splice blocking MOs (MO1: TAGGGTTTAGTCTTACCTTGGTCAC; MO2: TGATCTCTGATTGCTTACGTTCTTC) and one translation-blocking *atoh1b* MO (TCATTGCTTGTGTAGAAATGCATAT) were diluted in Danieau buffer containing 0.2% phenol red. 2–4 nL of *dll4* MO solution containing 4 ng oligonucleotide (MO1) and 3 ng oligonucleotide (MO2), and *atoh1b* MO solution containing 4 ng oligonucleotide were injected into the cell body of 1-cell stage embryos. Both *dll4* and *atoh1b* MO injected larvae were compared with larvae injected with a previously reported standard control MO (TACGGTTTACTCTTAGCTTCGTGAC)^104^ at 4 ng. Both control MO and *atoh1b*/*dll4* MO injected embryos were then raised at 28°C until 5 dpf when they were fixed, dehydrated, and stained using HCR for *best4*+ cells, enterochromaffin cells, and goblet cells. Around 10–15 embryos were stained and imaged for each MO injected.

### Drug inhibition of Notch signaling in zebrafish larvae

N-[N-(3,5–Difluorophenacetyl)-L-alanyl]-S-phenylglycine-t-butyl ester (DAPT, D5942, Sigma) is a γ-secretase inhibitor which was used to block Notch signaling. A 100X stock solution of 10mM DAPT in DMSO was prepared and stored at –20°C. Zebrafish larvae were incubated in 100 μM of DAPT in E3 media between 32-45 hours post fertilization (hpf), 48–72 hpf, and 72–120 hpf. Embryos incubated in 1% DMSO were used as controls.

### Heat shock treatments

For the misexpression of the Notch intracellular domain, we crossed Tg(*hsp:Gal4*)^kca4+/−^ and Tg(*UAS:myc-Notch1a-ICD*)*^kca3^* fish as described previously^90^, and heat-shocked at various developmental time windows – 48–72 hpf and 72–96 hpf. Each heat-shock session consisted of alternating incubations at 37 °C for 10 min and room temperature for 10 min, repeated over a total duration of 1 h. Within each developmental window, heat shock was performed three times—once at the beginning, once at the midpoint, and once during the final hour of the window. Following heat-shock treatment, larvae were maintained at 28 °C.

### CRISPR/Cas9 mediated whole gene deletion

Whole-gene deletion mutants for *meis1b^y730^* and *pbx3a^y731^* were generated using CRISPR/Cas9. For the *meis1b^y730^* mutation, four gRNAs, two targeting sequences spanning 63 bp upstream of the 5’ end of the gene (5’sgRNA1: 5’–CTGCTGAAACCCGGAAGTAGTGG–3’; 5’sgRNA2: 5’–TCAAGTTCCTGCTAAAGCAGGGG–3’) and spanning sequences 61 bp and 97 bp downstream from the 3’ end of the gene (3’sgRNA1: 5’–GTTTCTTCTCAAGGCCAAGACGG–3’; 3’sgRNA2: 5’–CAGGACGATGTGGCTCAGACAGG–3’) were designed using CRISPRscan and checked for off-target effects^105^. For the *pbx3a^y731^* mutant, four CRISPR/Cas9 guides were designed using the same parameters and tools, and targeted regions 24 bp and 71 bp upstream of the 5’ end (5’ sgRNA1: 5’–CCAGTATCAGAAGCAGTCTGTGT–3’; 5’ sgRNA2: 5’–CCAATAGTTAATATTGCACGCTA–3’) of the gene and 8 bp and 141 bp on the 3’ end (3’ sgRNA1: 5’–CTGCTGTGGGTTATCCGGTTTGG–3’; 3’ sgRNA2: 5’–TGGTGAGGTATCACCAGTTCTGG–3’).

The designed sgRNAs were obtained as crRNA and tracrRNA from IDT that was diluted to 100 μM with nuclease free duplex buffer and stored at –20°C. Stock Cas9 protein (61 μM) was diluted to 25 μM in Cas9 Dilution Buffer (20mM HEPES, pH = 7.5; 350 mM KCl, adjusted to pH = 7.5; 20% glycerol). crRNA and tracrRNA obtained from IDT were diluted to 100 μM with nuclease free duplex buffer and stored at –20°C. To form crRNA and tracrRNA duplexes, 5 μL of 100 μM crRNA and 5 μL of 100 μM tracrRNA were mixed and incubated at 95°C (5 mins), cooled to 25°C at a rate of 0.1°C per second, and maintained at 25°C for 5 mins. 0.8 μL of the crRNA/tracrRNA duplex was then mixed with 1:1:1 ratio of IDT duplex buffer, nuclease free water and 0.2% phenol red, and 1 μL of 25 μM prediluted Cas9 protein. The resultant mixture was then incubated at 37°C for 5 mins and was cooled to room temperature. The duplex mixtures containing crRNAs targeting the same gene were then pooled together and loaded onto an injection needle. For both mutants, one cell stage zebrafish embryos were injected with 20 μM Cas Protein (Integrated DNA Technologies) and each gRNA at a final concentration of 2 μM. Injected larvae were raised to adulthood and germline transmission of the mutation was confirmed using the following primer pairs: *meis1b* 5’Fwd1: 5’– CACACAGGCTTGTTCTTCACA –3’; *meis1b* 3’Rev1: 5’–TTGAGAATGGGTCCACGCTC–3’; *pbx3a* 5’Fwd1: 5’–AGTGTCCCTCGTTTGCACAA–3’; *pbx3a* 3’Rev1: 5’–CAGCAGCTCCTGAGGATGAG–3’.

F0 founders were then identified and outcrossed to wild-type fish to generate stable F1 generations. The F1 generations were then raised to adulthood and heterozygous carriers of the deletion were identified by genotyping PCR from fin clips. Experiments were performed with embryos derived from in-crosses of either F1 or F2 parents. Homozygous carriers of the deletion were identified using the following primer pairs in addition to the pairs mentioned above: *meis1b* 5’ Fwd2: 5’–TCCTCACACAGGCTTGTTCT–3’; *meis1b* 5’ Rev2: 5’– CCGTCTTGGCCTTGAGAAGA–3’; *pbx3a* 5’Fwd2: 5’–AGTGTCCCTCGTTTGCACAA–3’; *pbx3a* 5’Rev2: 5’– GGAGACGTGACTGATGACGG–3’. *meis1b^y730^* mutants further exhibited an elongated ear phenotype which was used to identify homozygous carriers. Larvae screened using the ear phenotype were further confirmed to be homozygotes using PCR genotyping when used for staining.

### Genotyping

For single-cell RNAseq, genotyping was performed by a combination of (a) micro fin clipping, (b) visual phenotype (for *meis1b^y730^* mutants), and (c) Zebrafish Embryo Genotyper (ZEG)^106^. For micro fin clipping, heterozygous F1 adults were in-crossed and F2 larvae were genotyped at 3 and 4 dpf by gently clipping a section of the caudal fin under a stereo microscope, distal to blood vessels, as previously described^107^. The fin clip sections were then lysed in 20 μL 50 mM NaOH solution at 95°C for 10 mins and subsequently cooled to 4°C in a thermocycler. 2 μL of Tris-HCl, pH = 7.4 was added to each sample, centrifuged and was used for setting up a PCR reaction to identify homozygous carriers and their WT siblings. For ZEG genotyping, a P20 pipette tip was cut using scissors, and with pipette set to 12 μL, was used to transfer a 3–4 dpf zebrafish larvae in E3 media + Tricaine on the circular holes on a ZEG slide. Once 24 animals had been transferred to the chip, the chip was loaded onto the platform of the base unit, the evaporation cover was placed over the chip, and the vibration motor was run at 1.8 volts for 7.5 minutes. Then using a P10 pipette, the liquid from each circular chamber was recovered without touching the embryo with the pipette tip. This liquid containing cells and genetic material of each larva was directly used for genotyping PCR reactions. The larvae were then transferred from the chip to a 24-well plate with blue water (5 mM NaCl, 0.17 mM KCl, 0.33 mM CaCl2, 0.33 mM MgSO4, 0.1% methylene blue) using a disposable 2ml transfer pipette. Blue water levels for each embryo in the 24 well plate were maintained at > 250 μL. *meis1b^y730^* wild-type siblings were identified by micro fin clipping, *meis1b^y730^* homozygous mutants were identified by visual phenotype, and *pbx3a^y731^* mutants were identified by both micro fin clipping and ZEG.

### Single-cell RNAseq

#### Dissociation of dissected zebrafish intestines into cell suspensions

Fertilized eggs were collected from *meis1b^y730^* and *pbx3a^y731^* mutant in-crosses and then reared in blue water (5 mM NaCl, 0.17 mM KCl, 0.33 mM CaCl2, 0.33 mM MgSO4, 0.1% methylene blue) at 28°C until they reached 6 dpf. Between 3–5 dpf, larvae were screened for the relevant phenotype or genotyped to distinguish homozygotes from their WT siblings, as described above. At 6 dpf, ∼80–100 intestines per sample were dissected by a team of 3–4 individuals, who alternated between WT and mutant samples to minimize dissection-related variability and prevent bias toward specific cell types. Dissected intestinal tissues were collected in 400 µL of sterile Ringer’s solution (116 mM NaCl, 2.6 mM KCl, 5 mM HEPES, pH 7.0) in 1.5 mL Eppendorf tubes. For each scRNA-seq experiment, protease solution (0.25% trypsin, 1 mM EDTA in 1× PBS, pH 8.0; Sigma-Aldrich T4549) was prepared fresh, and collagenase P (100 mg/mL in Hank’s Balanced Salt Solution; Sigma-Aldrich H9269 and 11213865001) was thawed on ice. For each sample, 1.2 mL of protease solution was added to a well of a 24-well plate, which was then incubated at 28 °C to equilibrate. The Ringer’s solution from each dissected sample (WT and mutant) was reduced to 100 µL, and the intestines were gently abraded by pipetting five times with a P200 set to 80 µL. Samples were rinsed by adding 1 mL fresh Ringer’s solution, allowing tissue to settle, and removing all but 100 µL. The intestines, along with the remaining Ringer’s, were then transferred into the equilibrated protease solution, followed by addition of 30 µL collagenase P per well. After swirling to mix (final volume 1.33 mL per well), plates were incubated at 28 °C and dissociation was monitored under a stereomicroscope. Every 5 minutes, samples were pipetted slowly 20 times with a P1000. After 10 minutes, each sample was passed once through a P200 by transferring from one well to another. Digestion was stopped after 20–25 minutes by adding 270 µL of 6× STOP solution (30% calf serum, 6 mM CaCl₂ in 1× PBS) and gently swirling (final volume 1.5 mL). Some small tissue fragments typically remained, but over-digestion usually reduced cell viability significantly and was avoided. Cell suspensions were then passed through a 40 µm cell strainer into 1.5 mL tubes, and each tube was flicked vigorously 20 times.

Cells were spun down at 300 x g for 3 minutes at 4°C. Supernatant was removed (leaving ∼ 100 μL) and cells were resuspended in 1 mL chilled DMEM/F12 (0% BSA; Gibco 12500062). Cells were washed by spinning at 300 x g for 3 minutes at 4°C, removing supernatant, and resuspending cells in 1 mL chilled DMEM/F12 (0% BSA). The single cell suspension was then filtered once again through a 40 µm cell strainer into clean tube, centrifuged at 300 x g for 3 minutes at 4°C. Supernatant was removed leaving ∼50 μL in which the cell suspension was resuspended.

#### Collection of single-cell transcriptomes

Droplet emulsions of single cell capture were generated using the 10X Genomics Chromium controller with Single Cell 3’ v3.1 consumable reagents for *pbx3a^y731^* mutant samples, and the 10X Genomics Chromium GEM–X platform with Single Cell 3’ v4 reagents for the *meis1b^y730^* mutant samples. Single-cell suspensions were stained with acridine orange / propidium iodide (10 μL of cells + 1 μL of Logos F23001 Acridine Orange/Propidium Iodide Stain) and then examined and quantified on a Logos Luna FL automated cell counter. Samples with sufficient concentration, low multiplet rate (<5% to proceed, <4.2% on average), and high viability (>85% to proceed, >90% on average) were then loaded into the instrument. WT and mutant samples for each mutant were then loaded into two channels at normal concentrations (*pbx3a^y731^*: 18,000 cells loaded, targeting 10,000 cells recovered; *meis1b^y730^*: 15,000 –18,000 cells loaded, targeting 11,000 – 13,000 cells recovered). Downstream reactions were performed in Eppendorf MasterCycler x50s Thermal cyclers. 10–12 cycles were used for cDNA amplification, and the result was inspected using Agilent D5000 ScreenTape Tapestation Assay. The remaining steps were performed according to the 10X 3’ v3.1 and 3’ v4 protocols, with 8–12 cycles used for sample index PCR, based on the concentration of amplified cDNA in the previous step. Final libraries were evaluated using Agilent D5000 ScreenTape Tapestation Assay and quantitated using a ThermoFisher Qubit 4 with dsDNA High Sensitivity (HS) reagents. Average fragment length from the Bioanalyzer and concentrations obtained from the Qubit were used to pool the WT and mutant libraries in equimolar concentrations. Libraries were then sequenced on a Novaseq 6000 Sequencing System (Illumina) using SP100-cycle full flowcells with 28 bases for Read 1, 10 bases for i7 index, 10 bases for i5 index, and 90 bases for Read 2. PhiX control library was spiked in at 1%. Reads obtained from these runs were used for downstream analyses.

#### Alignment of sequencing data

Alignment of sequencing reads and processing into a digital gene expression matrix was performed using Cell Ranger version 9.0.1, including the aligner STAR version 2.5.1b, with standard parameters. The data was aligned and quantitated against a custom reference index using Ensembl GRCz11 release 99 (January 2020) genomic FASTA, and using the Lawson Lab Zebrafish Transcriptome Annotation version 4.3.2, published in Lawson et al. eLife 2020^108^, available from https://www.umassmed.edu/lawson-lab/reagents/zebrafish-transcriptome/. Some light reformatting and reordering of entries was performed to make annotations consistent with Cell Ranger GTF formatting requirements, such as transcript/gene/exon grouping and matching the genomic reference ordering.

#### Normalization and quality control

Single cell transcriptomes for WT and mutant datasets were first processed separately using Seurat v4.1.0^109^ and R version 4.4.1. First, cells were scored (Seurat::PercentageFeatureSet) for their mitochondrial gene expression (using all genes beginning *mt-*) and ribosomal gene expression (using the genes: *rpl18a, rps16, rplp2l, rps13, rps17, rpl34, rpl13, rplp0, rpl36a, rpl12, rpl7a, rpl19, rps2, rps15a, rpl3, rpl27, rpl23, rps11, rps27a, rpl5b, rplp2, rps26l, rps10, rpl5a, rps28, rps8a, rpl7, rpl37, rpl24, rpl9, rps3a, rps6, rpl8, rpl31, rpl18, rps27.2, rps19, rps9, rpl28, rps7, rpl7l1, rps29, rpl6, rps8b, rpl10a, rpl13a, rpl39, rpl26, rps24, rps3, rpl4, rpl35a, rpl38, rplp1, rps27.1, rpl15, rps18, rpl30, rpl11, rpl14, rps5, rps21, rpl10, rps26, rps12, rpl35, rpl17, rpl23a, rps14, rpl29, rps15, rpl22, rps23, rps25, rpl21, rpl22l1, rpl36, rpl32, rps27l*). Cells from the *pbx3a^y731^* WT and mutant datasets were excluded if they showed either a low number of detected features and UMIs (≤200 genes or UMIs), unusually high complexity (≥3500 detected genes or ≥3000 UMIs), or elevated mitochondrial content (≥20%). The same filtering criteria was applied to the *meis1b^y730^* dataset, removing cells with ≤200 or ≥3500 detected genes/UMIs or with mitochondrial content ≥20%. Cells from all datasets were then log-normalized (Seurat::NormalizeData, normalization.method = “LogNormalize”, scale.factor = 10000) and scaled, regressing against mitochondrial and ribosomal gene expression (Seurat::ScaleData, vars.to.regress = c(“percent.mt”, “percent.ribo”)).

For each mutant, separate Seurat objects were created for the WT sibling and the homozygous mutants and subsequently combined using the base R “merge” function. For the *pbx3a^y731^*dataset, the *pbx3a*^−/−^ samples and the WT siblings were processed together on the same flowcell. In contrast, the *meis1b^y730^* single-cell libraries were generated across two experiments and sequenced on different flow cells due to technical failures in scRNAseq library generation. For visualization of the *meis1b^y730^* and *pbx3a^y731^* datasets, we identified the top 2000 variable genes (Seurat::FindVariableFeatures, selection.method = “vst”, nfeatures = 2000), performed PCA (Seurat::RunPCA), and identified significant PCs (Seurat::JackStraw, dims=100). WT and mutant samples for *meis1b^y730^* and *pbx3a^y731^* datasets were batch-corrected using harmony::RunHarmony (*pbx3a^y731^*: theta = 5, sigma = 0.1, lambda = 5; *meis1b^y730^*: theta = 2, sigma = 0.2, lambda = 2). Based on an elbow plot, 30 PCs were selected to use downstream. A Uniform Manifold Approximation and Projection (UMAP) was calculated using 50 nearest neighbors and the 30 most significant PCs (Seurat::RunUMAP, n.neighbors = 50). Leiden clustering was performed at multiple resolutions (Seurat::FindClusters, algorithm=4, resolution = c(0.75, 1, 2, and 3), n.start = 50, random.seed = 17). Marker genes were calculated for each cluster and used to manually curate and refine cluster identities. Clusters were annotated based on expression of markers identified in Daniocell. Differentially expressed genes between WT and mutant datasets in *meis1b^y730^* and *pbx3a^y731^* datasets were calculated using the command Seurat::FindAllMarkers(test.use = “roc”/”wilcox”, min.pct = 0.1, logfc.threshold = 0.1) applied to cells from each cluster individually. Results from the *best4*+ cell and goblet cell clusters are presented.

### Diffusion map analysis on *meis1b^y730^* mutants

Since *best4*+ cells are absent in *meis1b^y730^* mutants, we wanted to analyze what *best4*+ cell precursor transcriptional state can still be detected in the *meis1b^y730^* mutants. For that purpose, we extracted single-cell transcriptomes that represent secretory derivatives: secretory progenitor cells, goblet cells, enteroendocrine cells (including enterochromaffin cells), *best4*+ cells, and tuft-like cells. We then used the 30 most significant PCA embeddings to create a diffusion map using the R package “destiny” with 30 nearest neighbors (destiny::DiffusionMap, k = 30). Diffusion maps are a dimensionality reduction technique that reveals the underlying structure and continuum of the dataset that is useful for visualizing gradual transitions between cell states. Using the diffusion map, we tested whether the *meis1b^y730^* mutation (a) blocks secretory progenitors from progressing to *best4*+ cells or resulted in partial/incomplete *best4*+ cell differentiation by identifying any cells whose gene expression profiles had begun to shift toward that of *best4*+ cells.

We measured the Euclidean distance in DC space for each WT and *meis1b^y730^* homozygous mutant secretory progenitor to the centroid of the WT *best4*+ cluster in diffusion map space (diffusion components 1 and 2). We then selected the top 25% of WT and *meis1b*^−/−^ secretory progenitors that were closest to the centroid of the *best4*+ cell cluster. These progenitors were considered as the WT and *meis1b*^−/−^ secretory progenitors biased towards *best4*+ cells. Differential gene expression analysis was performed between these WT and *meis1b*^−/−^ secretory progenitors and *best4*+ cells using the command Seurat::FindMarkers(test.use = “roc”/”wilcox”, min.pct = 0.1, logfc.threshold = 0.25). Top differentially expressed markers were represented on a heatmap using the “pheatmap” package in R and on a volcano plot.

### Fluorescence Intensity Quantification

The values for the boxplot in Fig. 6O were obtained from confocal micrographs of *st6galnac1.1* HCR staining that were analyzed using Fiji (Image J) software to quantify fluorescence intensity differences between the WT and *pbx3a^y731^* samples. Images were converted to 8-bit grayscale and background-subtracted using a rolling ball radius of 100 pixels. Regions of interest (ROIs) were manually drawn around fluorescent cell populations in the distal intestine using the freehand selection tool, with 8 ROIs analyzed per genotype. For each ROI, the integrated density and area were measured, and background fluorescence was determined from cell-free regions outside the animal in the same image. Corrected total cell fluorescence (CTCF) was calculated using the formula: CTCF = Integrated density – (Area x Mean Background fluorescence). Statistical significance was assessed using an unpaired t-test with p < 0.05 considered significant.

### Gavage

6–7 dpf larvae were gavaged with ∼4-6 nL of 1% phenol red, 0.5% m-cresol, Alexa Fluor 647 dextran (ThermoFisher Cat# D22914), mCherry (Abcam, Cat# ab199750), or BODIPY™ FL C16 (ThermoFisher, Cat# D3821). Zebrafish larvae were briefly anesthetized with MESAB and then immobilized on a petri dish with 3% (wt/vol) methylcellulose (ThermoFisher, Cat# 900467-5) with the head tilted at a 45° angle to allow insertion of the needle into the mouth. Then, the oral microgavage technique was performed as previously described^110^, with a needle which passes through the mouth and esophagus to directly inject the injection mixture into the gut lumen of zebrafish larvae.

### Lipid and Protein uptake assays

To understand if *best4*+ cells uptake proteins and lipids, we microgavaged larval 7 dpf zebrafish with mCherry (Abcam, Cat# ab199750) or BODIPY™ FL C16 (ThermoFisher, Cat# D3821). 5 μL of 0.5 mg/mL BODIPY™ FL C16 was air dried and dissolved in 0.5 μL 100% EtOH and 4.5 μL 1X Danieau buffer (58 mM NaCl, 0.7 mM KCl, 0.4 mM MgSO_4_, 0.6 mM Ca(NO_3_)_2_, 5 mM HEPES, pH 7.6). mCherry protein aliquots were thawed and used at a concentration of 1 mg/mL. Larvae were gavaged with 4 nL of the 1 mg/mL mCherry protein and 0.5 mg/mL BODIPY™ FL C16 mixtures and imaged after 1 hour on a Nikon Eclipse Ti2 inverted microscope with a Nikon DS-R*i*2 camera using a 40X long working distance water/oil objective with Galvano settings. The gut was imaged with tile acquisitions that were later stitched together using the Nikon Elements Software.

### Luminal pH measurements

Larvae were microgavaged with pH indicators such as 1% phenol red (Sigma Cat# 143-74-8) and 0.5% m-cresol (ThermoFisher Cat# AAB2433806) mixed with acidic or neutral pH solutions. Acidic (pH 4) and neutral (pH 7) pH solutions were prepared by adding 1N HCl to D-PBS, while monitoring pH using a pH meter before mixing with pH indicators (phenol red and m-cresol). Microgavaged larval zebrafish guts were imaged on a Leica S9i stereomicroscope, which includes a color camera. D-PBS solutions of known pH were imaged within an Eppendorf tube to estimate the colors associated with each pH. The color of the bulb and pH standards were estimated using the ImageJ software plugin “RGB Measure” on an ROI of 150 pixels x 75 pixels. Another ROI of the same size was used outside the animal to quantify the background color for correction.

To quantify pH indicator colors as “hue” for graphing and analysis, first background correction was performed by linearly shifting RGB values of each image so that the RGB values of the background were set to the mean of all background RGB from an individual experiment. Then, RGB values were converted to Hue-Chroma-Luminance using farver::convert_colour. Since hue is circular, it was shifted by 180° so that the 0/360° boundary was not within the color range of the pH indicators used. Then, a fourth degree polynomial was fit to describe the Hue-Chroma relationship and Hue-Luminance relationship. There was a strong relationship, indicating that a single value (Hue) could be used to accurately describe the color of each animal, as expected, since the colors an individual pH indicator solution can adopt are limited. The hue value of each animal was then plotted over time, with the 95% confidence interval indicated.

### Intracellular pH measurements

pHluorin2 cloned into a middle-entry vector was obtained from Addgene #73794^98^. A stable line Tg(*best4:pHluorin2, cryaa:eGFP*)^y729^ expressing pHluorin2 under the control of the *best4* promoter was used for quantifying *best4*+ cells. Tg(*best4:pHluorin2, cryaa:eGFP*)^y729^ animals (N = 6) were mounted in 3% (w/v) methylcellulose at 5 and 6 dpf and cells in the anterior and posterior gut were imaged on a Nikon Eclipse Ti2 inverted microscope with a Nikon DS-R*i*2 camera using a 40X long working distance water/oil objective with Resonant settings at 4.17x magnification. A 20–50 section stack consisting of 1 µm z-slices was collected from the anterior and posterior intestinal regions. pHluorin2 fluorescence was acquired using both the 405 and 488 nm excitation lasers and a 525/50 emission filter. 5 µm z-stacks for individual cells across a ROI of 12 pixels x 12 pixels were used for image analysis. Background correction was performed by averaging background values per channel and then subtracting these values from the mean gray values of pHluorin2^+^ cells. Fluorescence values obtained at the 405 nm and 488 nm excitations were measured and then ratioed. 10–12 cells from the anterior intestine, and 3–6 cells from the posterior intestine were analyzed per animal.

### Quantification and Statistical Analysis

For all graphs, error bars report mean ± S.E.M. All graphs in Figures 3M, 3Q, 4D, 4G, 4J, 5J, 6O, S6J, S6N, S6R, S7E, S7H were plotted with ggplot2::geom_boxplot^111^ in R where the bar represents the median, the lower and upper hinges correspond to the first and third quartiles (the 25th and 75th percentiles), and the whiskers extend from the hinge to the value most distant from the mean and no further than 1.5 * IQR from the hinge (where IQR is the inter-quartile range, or distance between the first and third quartiles). Graphs in Figure 7H and S11E were plotted using GraphPad Prism 10. Heatmaps were generated with ggplot2::geom_tile() in Fig. 1, and Fig. S1 and using the “pheatmap” function in Fig. 5 and Fig. S9. All computational analysis and statistical tests were performed in either R or GraphPad Prism v10 (GraphPad software, San Diego, CA). The rest of the data processing steps are described in their respective methods sections. Figure legends describe the details of the data plotted including what statistical tests were performed, significance, and sample sizes.

