## Supplementary Figures S1 - S11 for "Developmental regulation of intestinal *best4*+ cells"

### Supplementary Material for
### Developmental regulation of intestinal *best4*+ cells

Abhinav Sur[1,‡], Ella X. Segal[1,‡], Michael P. Nunneley[1], Jason W. Sinclair[1], Morgan Kathleen Prochaska[1], Louis E. Dye[2], Yalan Wu[1], Liezhen Fu[3], Yun-Bo Shi[3], James Iben[4], Benjamin Feldman[1], Jeffrey A. Farrell[1,*]

[1] Division of Developmental Biology, *Eunice Kennedy Shriver* National Institute of Child Health and Human Development, NIH, Bethesda, MD 20892, USA.

[2] Microscopy and Imaging Core, *Eunice Kennedy Shriver* National Institute of Child Health and Human Development, NIH, Bethesda, MD 20892, USA.

[3] Section on Molecular Morphogenesis, *Eunice Kennedy Shriver* National Institute of Child Health and Human Development, NIH, Bethesda, MD 20892, USA.

[4] Molecular Genomics Core, *Eunice Kennedy Shriver* National Institute of Child Health and Human Development, NIH, Bethesda, MD 20892, USA.

‡ These authors contributed equally.

**<u>List of Supplementary Materials</u>**

**Supplementary Figures S1 – S11 (this PDF)**

**Video S1:** Timelapse confocal imaging of 84 hpf Tg(*best4:mApple*) zebrafish showing *best4*+ cells (magenta) in the intestinal bulb extending and retracting cytoplasmic protrusions in many orientations. Related to Fig. 2B–C.

**Video S2**: Live confocal micrograph of 96 hpf *atoh1b*$^{mGL}$; Tg(*best4:mApple*) zebrafish showing *atoh1b*$^{mGL}$ + cells expressing Tg(*best4:mApple*) throughout the length of the gut. Some *atoh1b*$^{mGL}$+; Tg(*best4:mApple*)+ cells can also be seen turning off *atoh1b*$^{mGL}$ expression. Related to Fig. 3C–F.

**Video S3**: Timelapse confocal imaging of 76 hpf *atoh1b*$^{mGL}$; Tg(*best4:mApple*) zebrafish showing *atoh1b*$^{mGL}$ + cells turning on Tg(*best4:mApple*) expression indicating *best4*+ cells descend from *atoh1b* secretory progenitors. Related to Fig. 3D.

**Table S1**: HCR probes used in this study to profile secretory progenitors, *best4*+ cells, enterochromaffin cells, goblet cells, and tuft cells under different conditions.

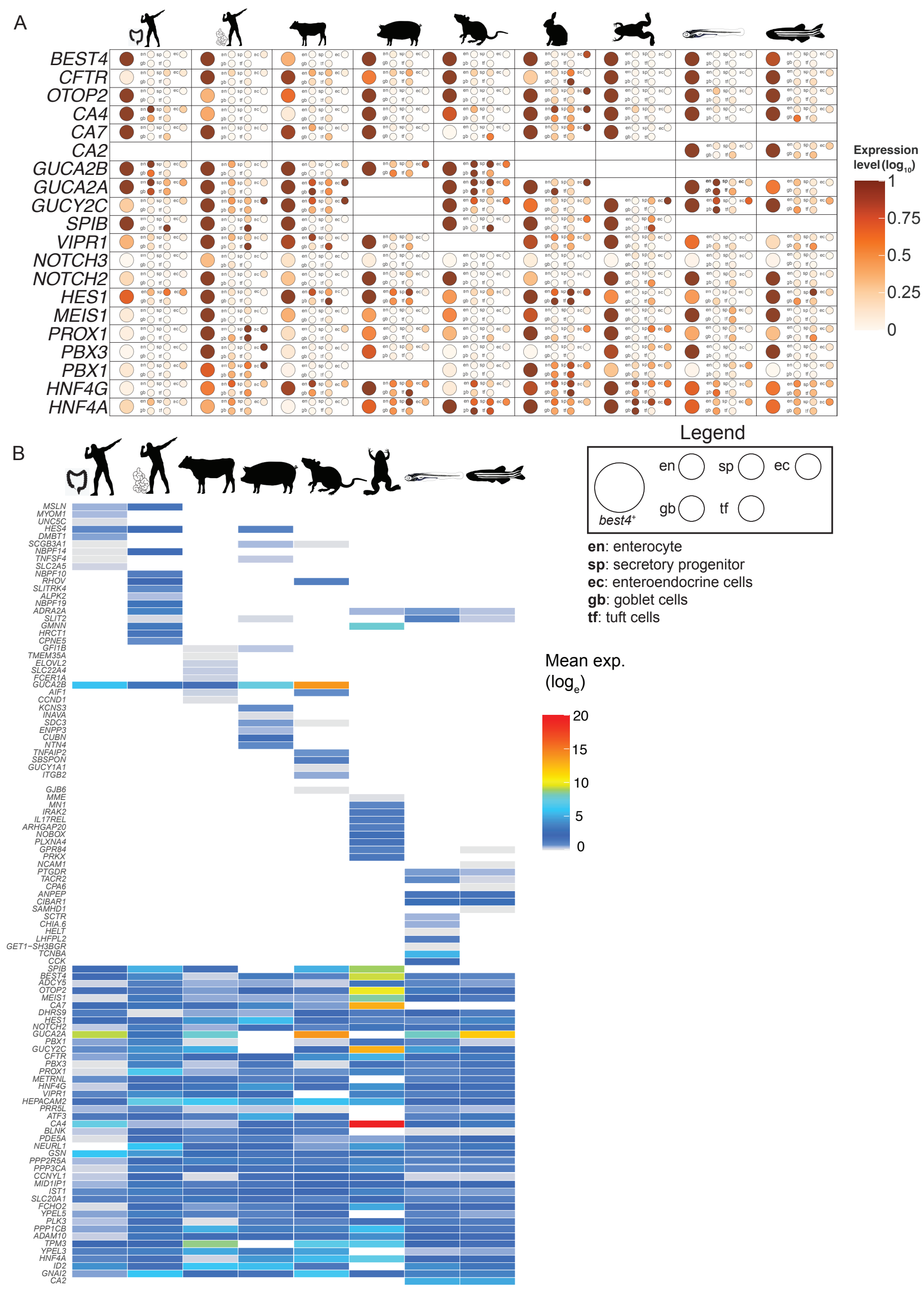


**Figure S1**: **Expression of conserved *best4*+ cell-specific genes across zebrafish intestinal cell types and species.** Related to Figure 1.

(**A**) Dot plot of mean expression ($log_{10}$) of the most highly conserved *best4*+ cell-specific genes in *best4*+ cells (big circle) and other IECs (en: enterocyte, sp: secretory progenitors, ec: enterochromaffin cells, gb: goblet cells, tf: tuft/tuft-like cells) across animals.

(**B**) Heatmap showing most differentially expressed genes (i.e. the most highly expressed markers) of *best4*+ cells in each analyzed species. Organized with species-specific signatures on top, followed shared markers below.

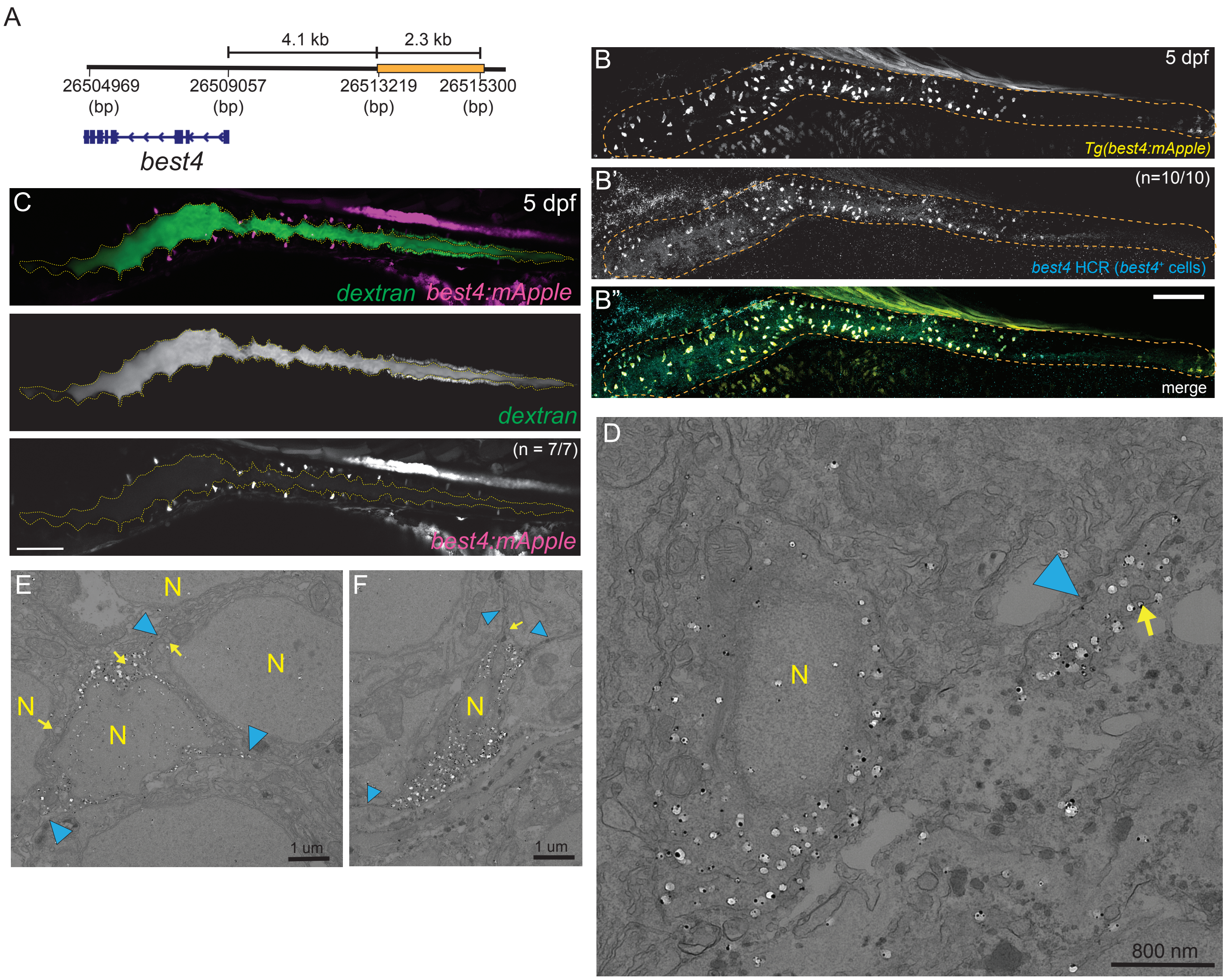


**Figure S2**: **Live imaging and electron microscopy using a live *best4*⁺ cell reporter line reveals dynamic protrusions of *best4*⁺ cells.** Related to Figure 2.

(**A**) Cartoon showing the 4.1 kb upstream genomic region (orange) that was used to drive reporter gene expression in *best4*⁺ cells.

(**B–B"**) Validation of Tg(*best4:mApple*) expression in 5 dpf zebrafish larvae using HCR RNA *in situ* hybridization against *best4* mRNA and immunofluorescence against mApple. Scale bar: 100 um

(**C**) Confocal images of a 5 dpf Tg(*best4:mApple*) zebrafish gut gavaged with fluorescent dextran (green) showing *best4*⁺ cells extend dynamic projections in all regions of the zebrafish gut. Scale bar: 100 um.

(**D–F**) Immuno-electron microscopy showing *best4*+ cells detected using a GFP antibody in a *Tg(best4:eGFP)* background. Arrowheads indicate cytoplasmic protrusions; arrows denote subapical vesicles. Scale bar: as indicated in each panel.

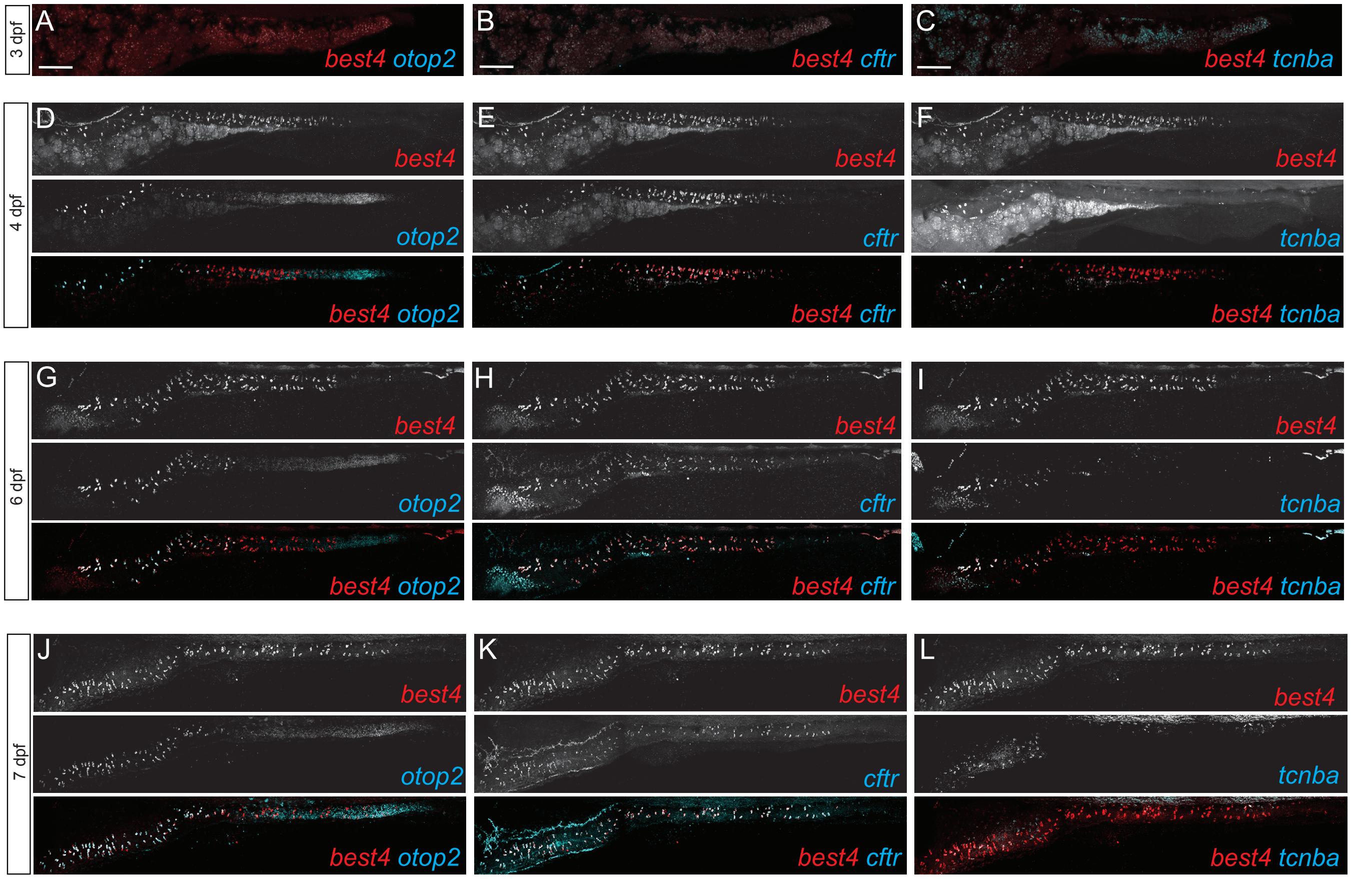


**Figure S3: Gene expression regionalization along anterior–posterior axis within *best4*⁺ cells remains consistent across developmental stages.** Related to Figure 2.

(**A–L**) HCR RNA *in situ* hybridization showing gene co-expression of *best4*+ cell markers in 3 dpf (prior to the onset of *best4*+ cell gene expression) (A–C), 4 dpf (D–F), 6 dpf (G–I), and 7 dpf (J–L). RNA *in situ* hybridization of these markers in 5 dpf larvae shown in Figure 2K–M. Scale bar: 100 um.

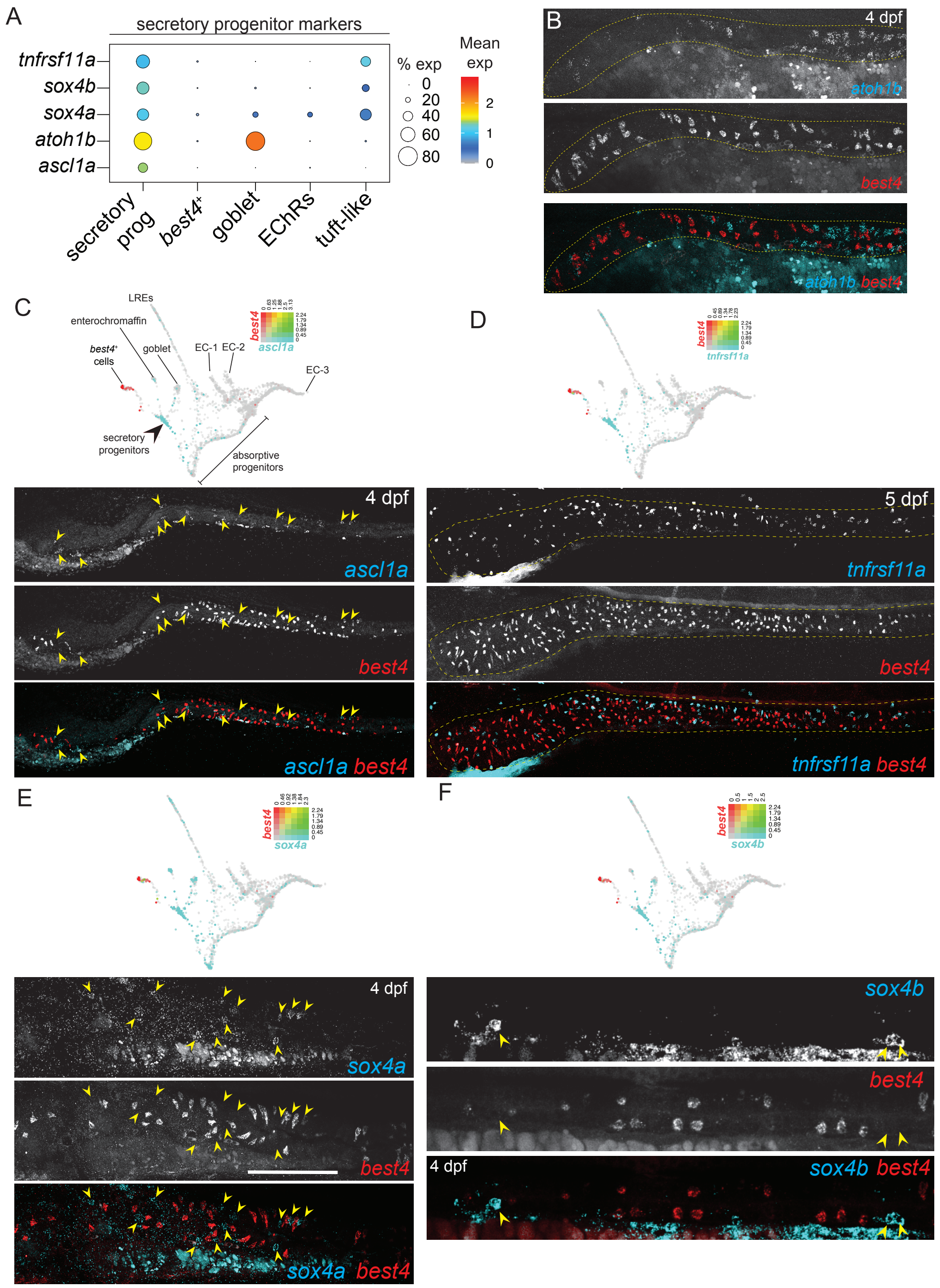


**Figure S4: Secretory progenitor marker gene expression does not overlap *best4*+ cells at the mRNA level**. Related to Figure 3. (**A**) Dot plot of secretory progenitor markers (Y-axis) identified in our previous study across secretory derivatives (X-axis).

(**B–F**) Lack of co-expression of secretory markers *ascl1a, tnfrsf11a, sox4a, and sox4b* (cyan) with *best4* (red) on URD-inferred transcriptional trajectory of intestinal epithelial cells (Sur et al 2023). HCR RNA *in situ* hybridization for *atoh1b* (**B**), *ascl1a* (**C**), *tnfrsf11a* (**D**), *sox4a* (**E**), and *sox4b* (**F**) confirms lack of co-expression of secretory progenitor markers and *best4*. Scale bar: 100 um.

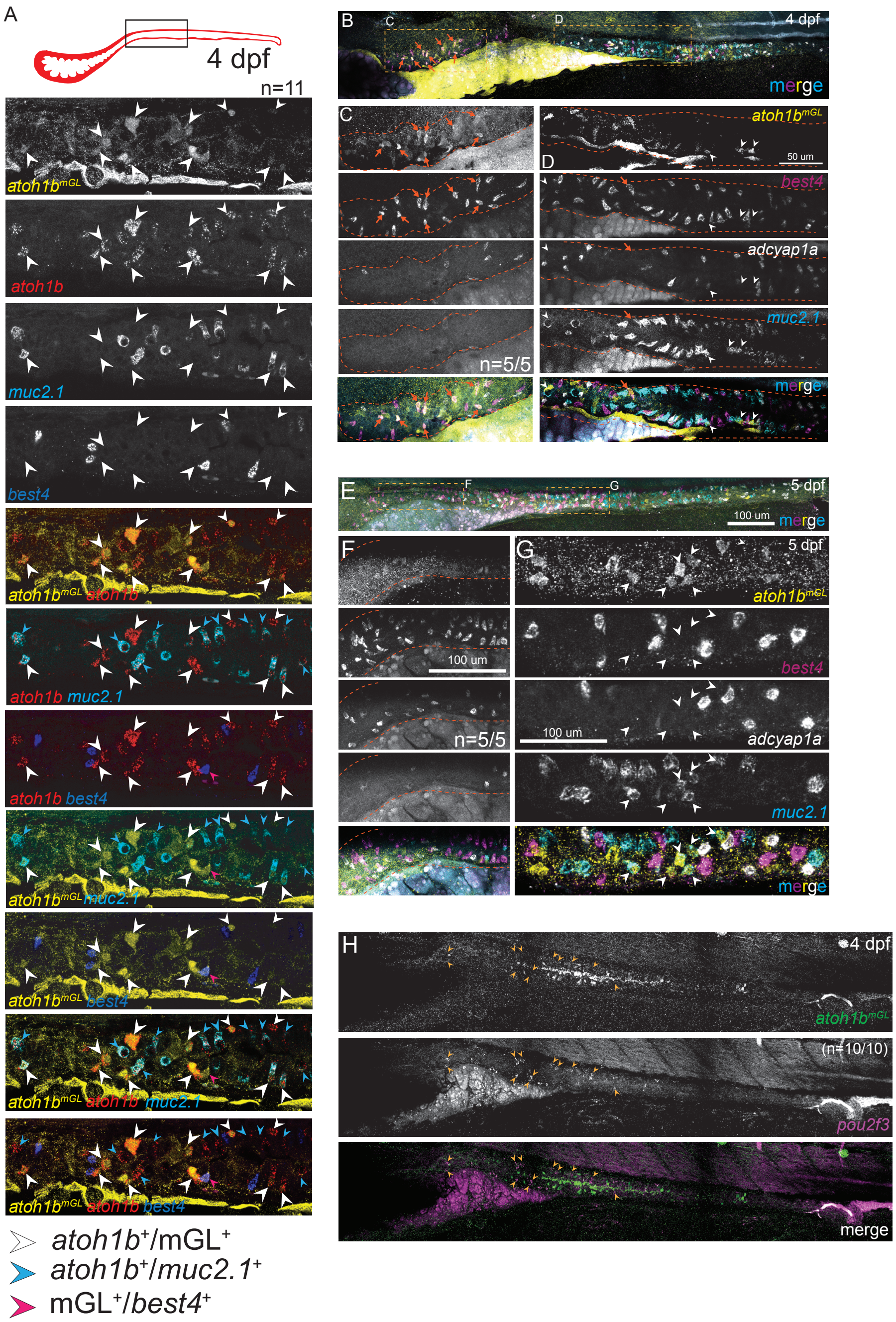


**Figure S5**: **Characterization of dynamic *atoh1b* expression and *atoh1b*$^{mGL}$ labeling of secretory progenitors and secretory derivatives**. Related to Figure 3.

(**A**) HCR RNA *in situ* hybridization of secretory master regulator *atoh1b* along with *best4*+ cell (*best4*), and goblet cell (*muc2.1*) markers on an immunofluorescent *atoh1b*$^{mGL}$ background. Most *atoh1b*+ cells are either *atoh1b*$^{mGL}$+ secretory progenitors (white arrowheads) or goblet cells (*muc2.1*+, cyan arrowheads). Most *atoh1b*$^{mGL}$+ cells were either *atoh1b*+ (white arrowheads) or *best4*+ (magenta arrowheads). Scale bar: 50 um.

(**B–G**) HCR RNA *in situ* hybridization of *best4*+ cell (*best4*), enterochromaffin cell (*adcyap1a*) and goblet cell (*muc2.1*) markers co-stained with *atoh1b*$^{mGL}$ immunofluorescence at 4 dpf (**B–D**), and 5 dpf (**E–G**). (**B, E**) Overview of intestine indicating anterior and posterior regions shown at high magnification in (**C–D, F–G**). *atoh1b* expression is dynamic and moves from the anterior to posterior of the intestine between 4 and 5 dpf. Red arrows indicate overlapping *atoh1b*$^{mGL}$ immunofluorescence signal with *best4*$^{+}$ cells (predominantly observed at 4 dpf) and white arrowheads denote *atoh1b*$^{mGL}$ immunofluorescent staining in *muc2.1*$^{+}$ goblet cells (predominantly observed at 5 dpf). (**H**) *atoh1b*$^{mGL}$ immunofluorescence co-stained with RNA *in situ* hybridization of tuft-like cell marker *pou2f3*. Arrowheads indicate *pou2f3*$^{+}$ tuft-like cells that are not marked by *mGL* immunofluorescence. Co-expressing cells were not observed.

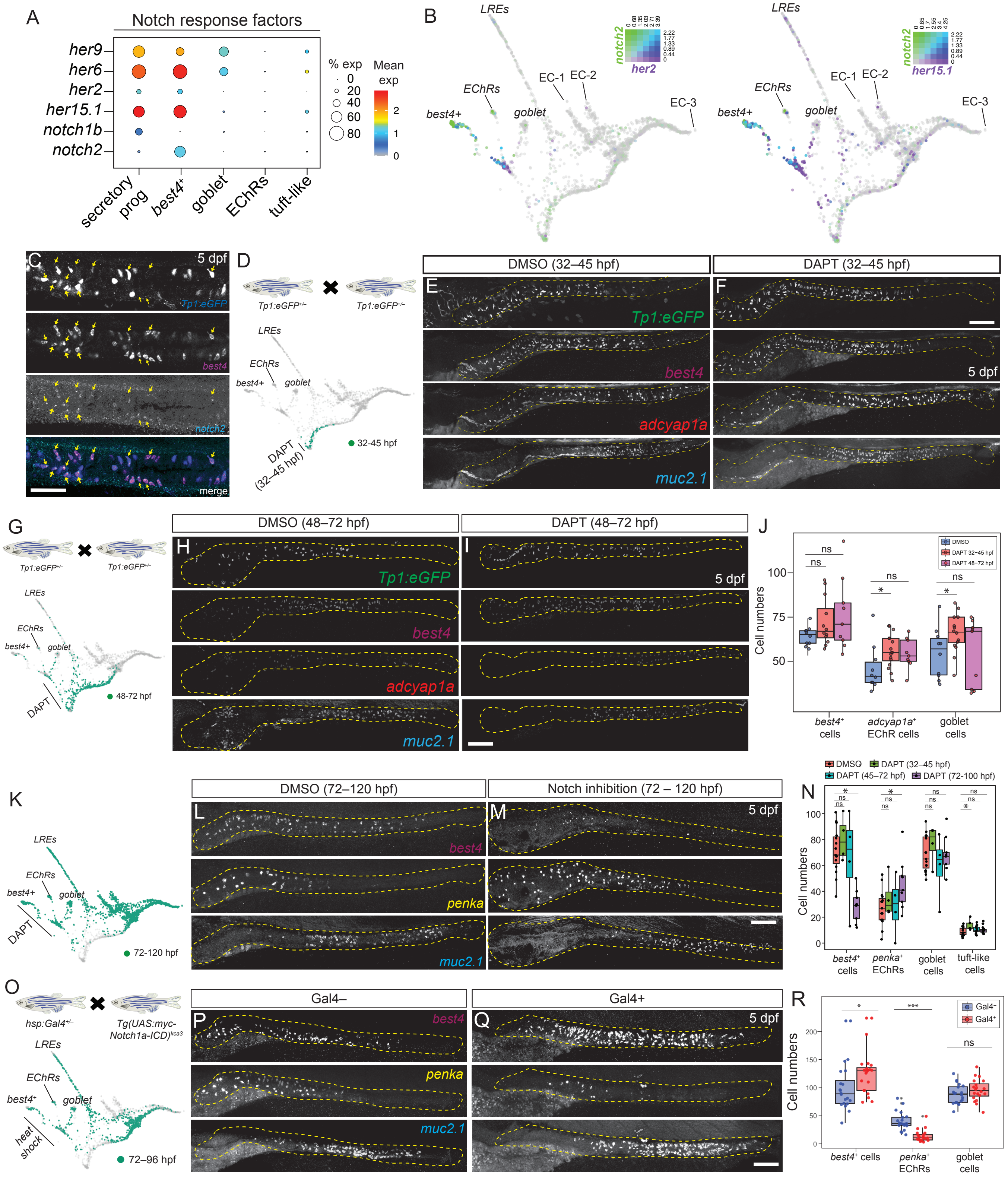


**Figure S6: Notch signaling between 72–120 hpf increases *best4*+ cell number and reduces enterochromaffin cell number.** Related to Figure 4.

(see next page for Legend)

**Figure S6: Notch signaling between 72–120 hpf increases *best4*⁺ cell number and reduces enterochromaffin cell number.** Related to Figure 4.

(**A**) Expression of Notch receptors and effectors (Her/Hes factors) across secretory derivatives.

(**B**) Co-expression of the Notch receptor *notch2* (green) and Hes5 Notch effectors (*her2*, *her15.1*, purple) on URD-inferred transcriptional trajectory of intestinal epithelial cells (Sur et al 2023).

(**C**) HCR RNA *in situ* hybridization of *best4*+ cells (*best4*) and *notch2* on a Tg(*Tp1-bglob:eGFP*) Notch reporter line immunofluorescence background. Yellow arrows indicate cells actively receiving Notch signaling (GFP+) that express both *notch2* and *best4* mRNA. Scale bar: 50 um.

(**D–N**) Alteration of cell type proportions following inhibition of Notch signaling at different developmental stages using DAPT, a γ-secretase inhibitor. (**D**, **G**, **K**) Experimental plan for each experiment, including indication of cells from perturbed developmental stages on URD-inferred transcriptional trajectory of intestinal epithelial cells (Sur et al 2023). (**E, F, H, I, L, M**) HCR RNA *in situ* hybridization at 5 dpf to mark *best4*+ cells (*best4*), enterochromaffin cells (*adcyap1:* **E, F, H, I**; *penka*: **L, M**) and goblet cells (*muc2.1*) with Tg(*Tp1:eGFP*) immunofluorescence (**E, F, H, I**) to demonstrate recovered Notch signaling after treatment with DMSO (**E, H, L**) or DAPT (**F, I, M**). (**J**) Quantification of number of *best4*⁺ cells, enterochromaffin cells, and goblet cells after Notch inhibition between 32–45 hpf and 48–72 hpf, presented in **D–I**. (**N**) Quantification of number of *best4*⁺ cells, anterior enterochromaffin cells (*penka*+), goblet cells, and tuft-like cells after Notch inhibition between 32–45 hpf, 45–72 hpf, and 72–120 hpf (shown in **K–M**). EChR: enterochromaffin. Scale bar: 100 um.

(**O–R**) Increased Notch signaling via heat shock of Tg(*hsp:Gal4*); Tg(UAS:myc-Notch1a-ICD) larvae. (**O**) Schematic of experimental plan, indicating cells from perturbed timepoints. (**P–Q**) HCR RNA *in situ* hybridization to mark *best4*+ cells (*best4*), enterochromaffin cells (*penka*) and goblet cells (*muc2.1*) in heat-shocked larvae and their Gal4– siblings. (**R**) Quantification of cell numbers after heat shock. Scale bar: 100 um.

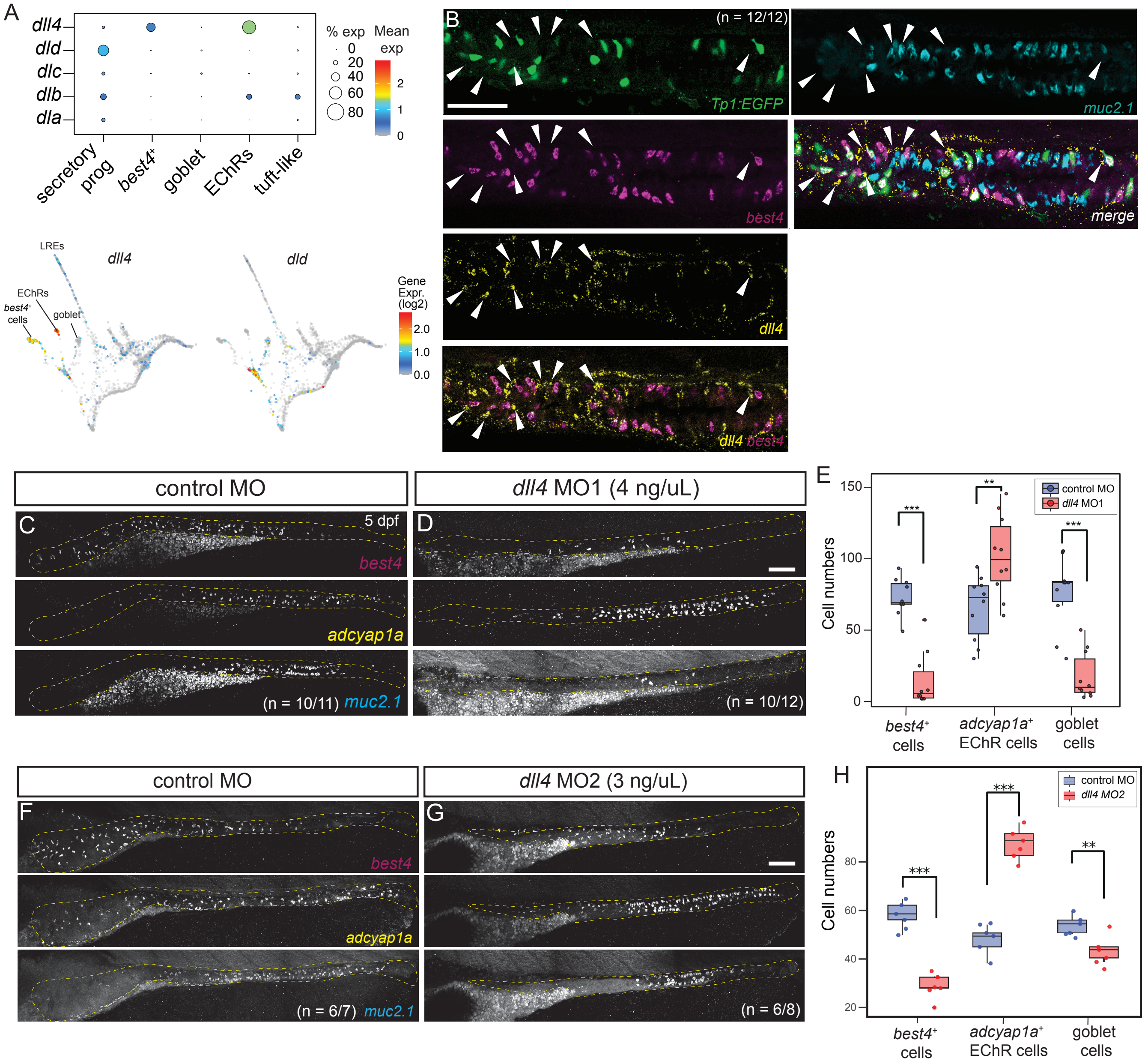


**Figure S7: *dll4* is a key Notch ligand required for *best4*+ cell specification.** Related to Figure 4.

(**A**) Gene expression of Notch ligands (i.e., Delta) across secretory derivatives as a dot plot and on URD-inferred transcriptional trajectory of intestinal epithelial cells (Sur et al 2023).

(**B**) RNA *in situ* hybridization of *dll4* (*delta-like 4*) along with *best4*+ cell (*best4*) and goblet cell (*muc2.1*) markers co-stained with Tg(*Tp1-bglob:eGFP*) Notch reporter immunofluorescence. Arrowheads indicate *dll4*+ cells that are adjacent to a Tg(*Tp1-bglob:eGFP*)+ or *best*4+ cell. Scale bar: 50 um.

(**C–H**) HCR RNA *in situ* staining in 5 dpf larvae of *best4*+ cell (*best4*), enterochromaffin cell (*adcyap1a*) and goblet cell (*muc2.1*) markers after injection of control MO (**C, F**), *dll4* MO1 (**D**), and *dll4* MO2 (**G**). Boxplots showing number of *best4*+ cells, enterochromaffin cells, and *muc2.1*+ goblet-cells after *dll4* MO1 (**E**) and *dll4* MO2 (**H**) mediated knock-down vs control MO injected larvae. Colors indicate control MO (blue) and morpholino injected animals (red). Scale bar: 100 um.

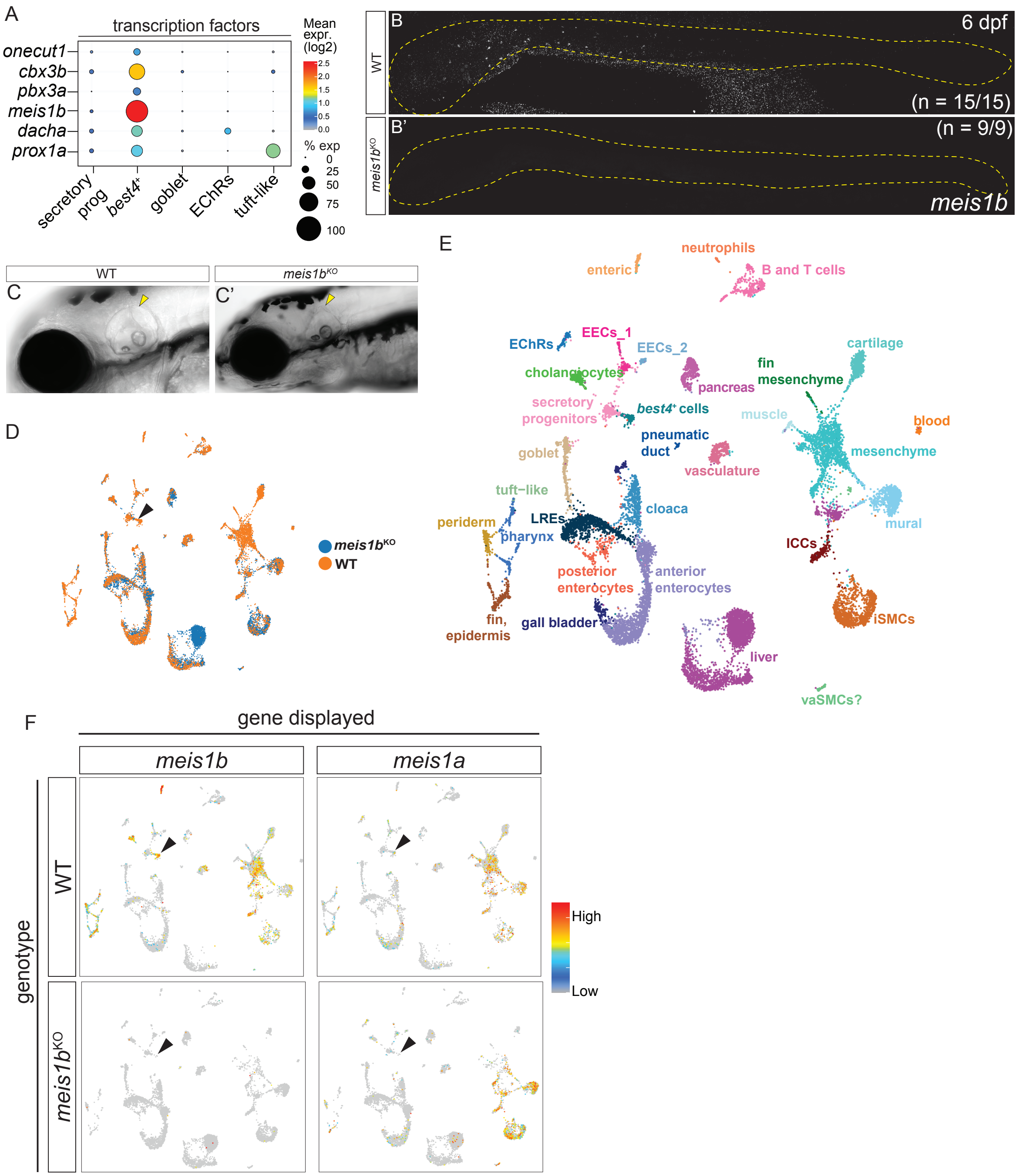


**Figure S8**: **Whole gene deletion of *meis1b* leads to a complete loss of *meis1b* transcript and *best4*$^+$ cells.** Related to Figure 5.

(**A**) Expression of key transcription factors specifically enriched in *best4*$^+$ cells among secretory derivatives.

(**B, B'**) HCR RNA *in situ* hybridization showing loss of *meis1b* transcript in *meis1b*$^{-/-}$ 6 dpf zebrafish larval intestine (**B'**) compared to wild-type siblings (**B**). *meis1b* signal is weak even in wild-type animals due to its low expression level.

(**C, C'**) DIC images of 5 dpf zebrafish larval heads showing a normal inner ear phenotype in wild-type larvae (**C**) and an elongated ear phenotype (**C'**; yellow arrowheads) in *meis1b*$^{-/-}$ larvae.

(**D–E**) UMAP projection of 6337 *meis1b*$^{+/+}$ cells and 8012 *meis1b*$^{-/-}$ cells dissociated from dissected zebrafish intestines at 6 dpf, colored by genotype (**D**) and cluster identity (**E**). The *best4*$^+$ cell cluster is shown using a black arrowhead.

(**F**) Expression (natural log) of *meis1b* and paralog *meis1a* in WT vs *meis1b*$^{-/-}$ dataset.

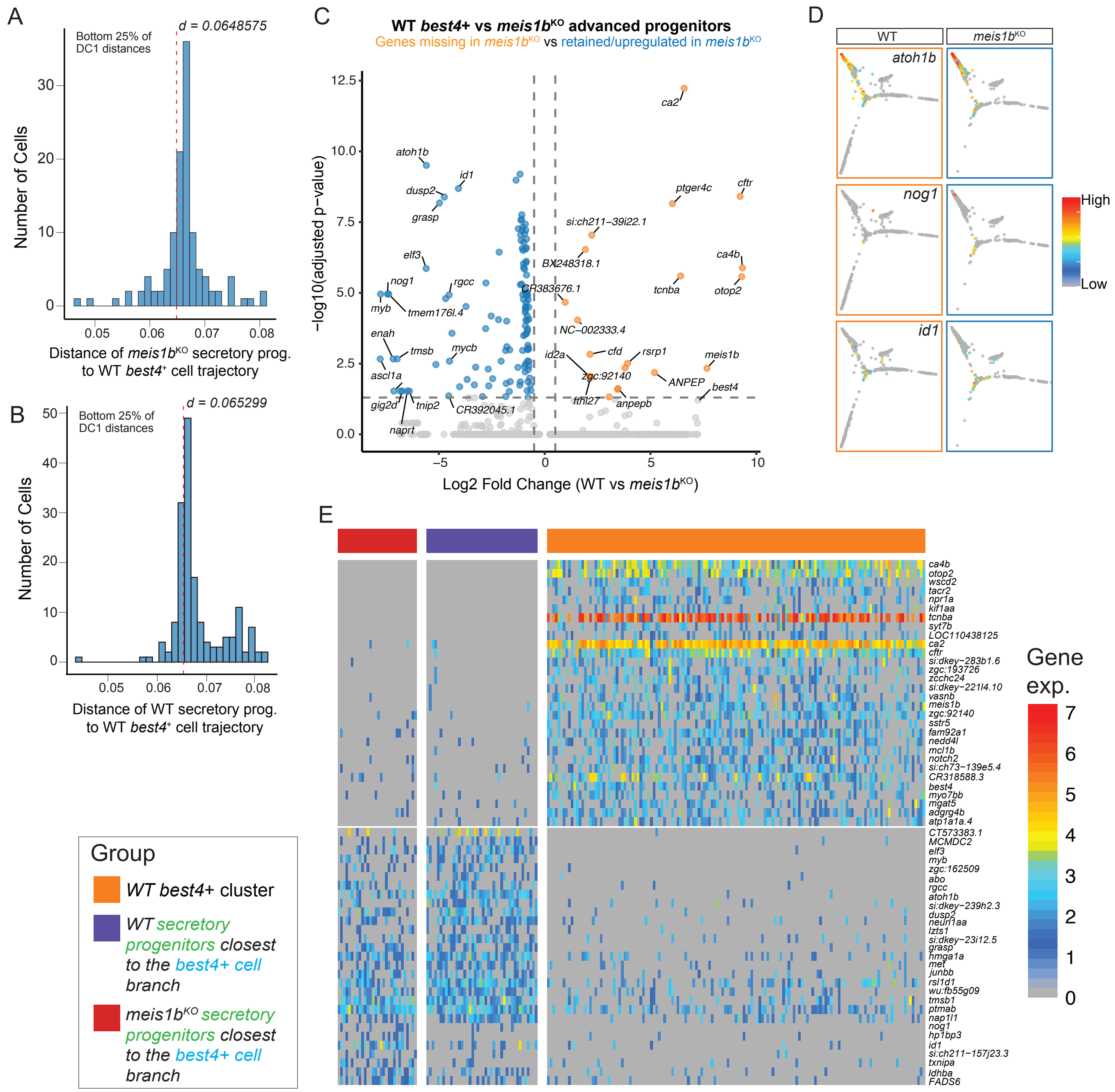


**Figure S9: *meis1b*–/– mutant secretory progenitors resemble WT *best4*+ cell biased secretory progenitors but don't turn on *best4*+ cell differentiation markers.** Related to Figure 5.

(**A–B**) Histogram showing the distribution of distances of *meis1b*–/– secretory progenitors (**A**) and wild-type secretory progenitors (**B**) to the *best4*+ cell cluster in diffusion component space. Red dotted line shows the 25% DC1 distance cutoff. Progenitors whose distances to the *best4*+ cell cluster was less than the red dotted lines were considered as secretory progenitors biased towards *best4*+ cells.

(**C**) Volcano plot of genes differentially expressed between WT *best4*+ cells (orange) and *meis1b*–/– *best4*+ cell biased secretory progenitors (blue). Select genes labeled on the plot.

(**D**) Diffusion map of secretory cell types colored by gene expression of selected differentially expressed genes from Fig. 5L.

(**E**) Grouped heatmap showing the top differentially expressed genes between the WT *best4*+ cell cluster, WT secretory progenitors biased towards *best4*+ cells, and *meis1b*–/– secretory progenitors biased towards *best4*+ cells. Color indicates natural log transformed expression level.

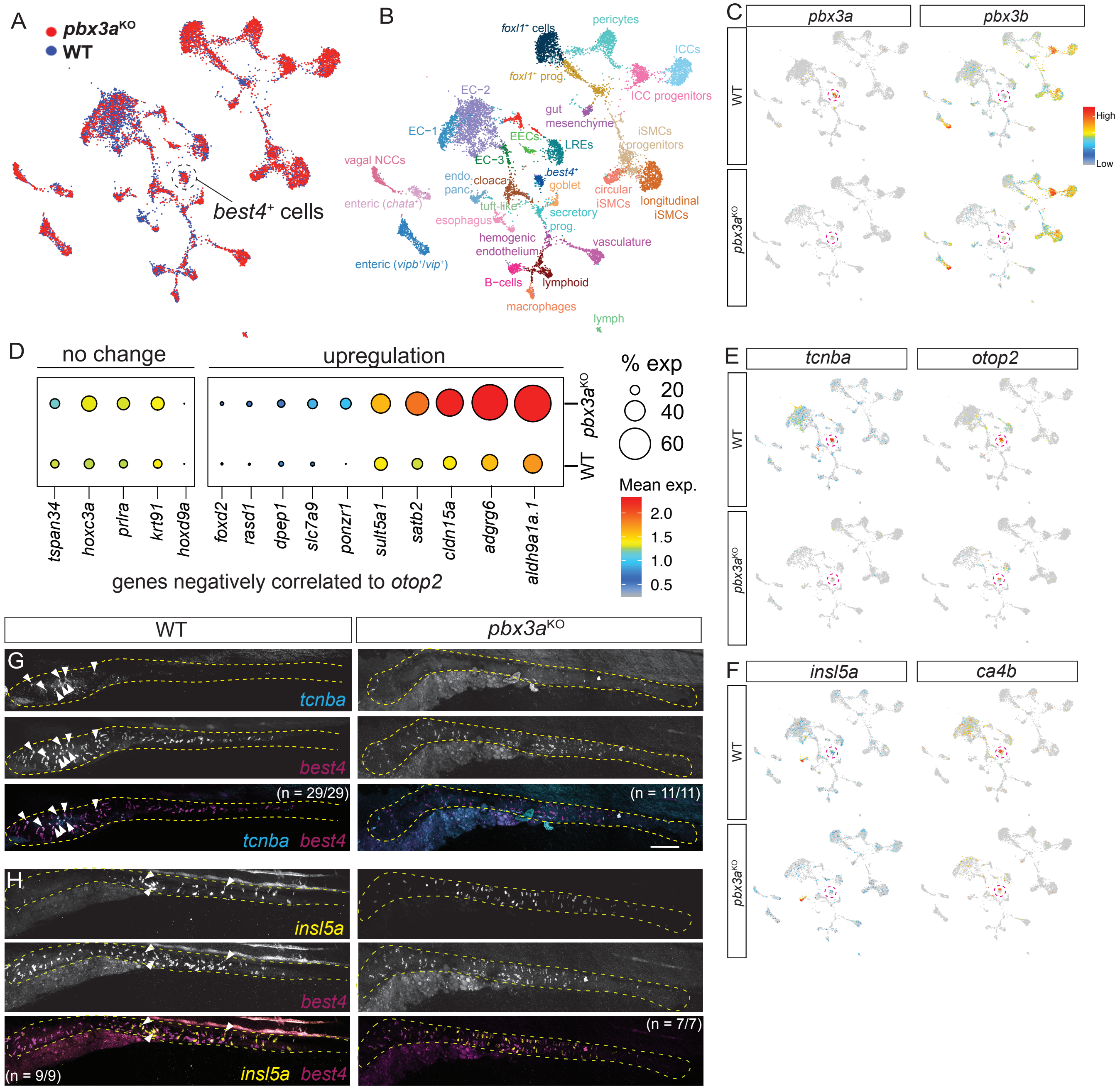


**Figure S10**: ***pbx3a$^{-/-}$* mutants exhibit lower expression of anteriorly regionalized genes and potential upregulation of putative posteriorly regionalized genes.** Related to Figure 6.

(**A, B**) UMAP projection of 6455 *pbx3a*$^{+/+}$ cells and 6099 *pbx3a*$^{-/-}$ single-cell transcriptomes isolated from dissected zebrafish intestines at 6 dpf colored by genotype (**A**) and cluster identity (**B**). *best4*$^{+}$ cell cluster indicated using a dotted circle.

(**C**) Expression (natural log) of *pbx3a* and paralog *pbx3b* in WT vs *pbx3a*$^{-/-}$ dataset. *best4*$^{+}$ cell cluster indicated using a dotted circle.

(**D**) Gene expression of *best4*+ cell markers negatively correlated to *otop2* (Fig. 2J), compared between *pbx3a* mutants and wild-type siblings. Expression is natural log transformed.

(**E–F**) Expression (natural log) of differentially expressed genes in the *pbx3a*$^{-/-}$ dataset compared to WT. *best4*$^{+}$ cell cluster indicated using a dotted red circle.

(**G–H**) HCR of putatively spatially heterogeneous genes (*tcnba* and *insl5a*) differentially expressed between *pbx3a*$^{-/-}$ mutants and WT siblings. Arrowheads indicate co-expressing cells.

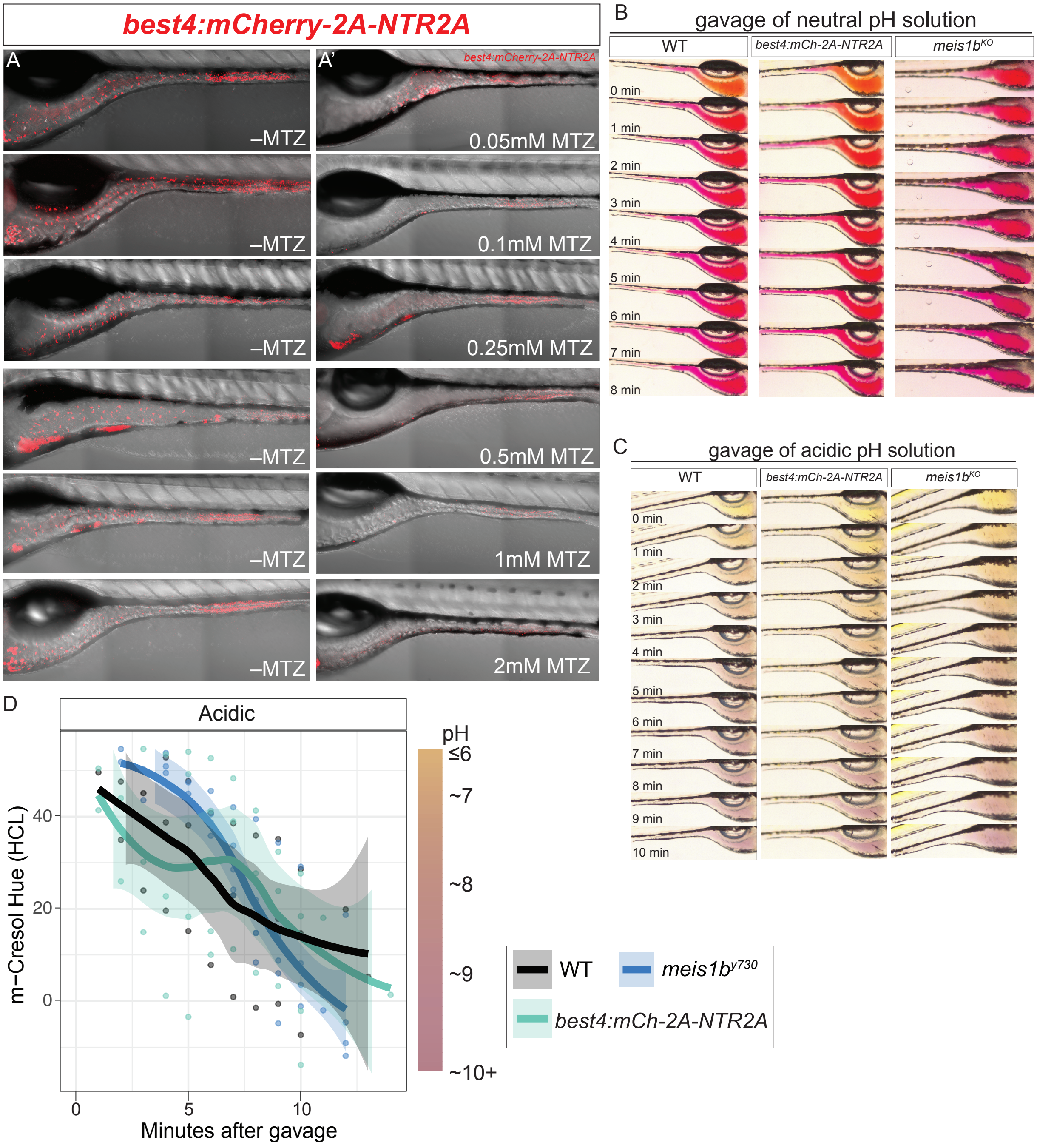


**Figure S11: Genetic ablation model that removes *best4*+ cells on command shows that larval intestines lacking *best4*+ cells can return to homeostatic pH following acidic challenge.** Related to Figure 7.

(**A**) Confocal imaging of Tg(*best4:mCherry-2A-NTR2A*) larval intestines (5 dpf) following treatment with different doses of metronidazole (MTZ) from 4.5 – 5 dpf. *best4*+ cells are effectively ablated with 0.1mM MTZ or higher.

(**B–C**) Images after microgavage of neutral (pH 7) 1% phenol red or acidic (pH 4) 0.5% m-cresol purple into wild-type, 0.1 mM metronidazole treated Tg(*best4:mCh-2A-NTR2A*), or *meis1b*$^{-/-}$ mutant larvae.

(**D**) Quantification of color changes after microgavage with acidic (pH 4) 0.5% m-cresol purple.
